# Bi-allelic *MCM10* mutations cause telomere shortening with immune dysfunction and cardiomyopathy

**DOI:** 10.1101/844498

**Authors:** Ryan M. Baxley, Wendy Leung, Megan M. Schmit, Jacob Peter Matson, Marissa K. Oram, Liangjun Wang, John Taylor, Lulu Yin, Jack Hedberg, Colette B. Rogers, Adam J. Harvey, Debashree Basu, Jenny C. Taylor, Alistair T. Pagnamenta, Helene Dreau, Jude Craft, Elizabeth Ormondroyd, Hugh Watkins, Eric A. Hendrickson, Emily M. Mace, Jordan S. Orange, Hideki Aihara, Grant S. Stewart, Edward Blair, Jeanette Gowen Cook, Anja-Katrin Bielinsky

## Abstract

Minichromosome maintenance protein 10 (Mcm10) is essential for eukaryotic DNA replication. Here, we describe compound heterozygous *MCM10* mutations in patients with distinctive but overlapping clinical phenotypes – natural killer (NK) cell deficiency (NKD) and restrictive cardiomyopathy (RCM) with hypoplasia of the spleen and thymus. To understand the mechanism of Mcm10-associated disease, we modeled these mutations in human cell lines. Mcm10 deficiency causes chronic replication stress that reduces cell viability due to increased genomic instability and telomere erosion. Our data suggest that loss of Mcm10 function constrains telomerase activity by accumulating abnormal replication fork structures enriched with single-stranded DNA. Terminally-arrested replication forks in Mcm10-deficient cells require endonucleolytic processing by Mus81, as *MCM10*:*MUS81* double mutants display decreased viability and accelerated telomere shortening. We propose that these bi-allelic mutations in *MCM10* predispose specific cardiac and immune cell lineages to prematurely arrest during differentiation, causing the clinical phenotypes in both NKD and CM patients.

## INTRODUCTION

The DNA replication program has evolved to ensure the fidelity of genome duplication. Conceptually, this program can be divided into origin licensing, origin firing and DNA synthesis. Origins are licensed by loading double hexamers of the core replicative helicase composed of minichromosome maintenance complex proteins 2 to 7 (Mcm2-7)^1, 2^. Origin firing requires helicase co-activators, including the go-ichi-ni-san (GINS) complex and cell division cycle protein 45 (Cdc45), to form the Cdc45:Mcm2-7:GINS (CMG) helicase^3, 4^. Upon firing, minichromosome maintenance protein 10 (Mcm10) is essential for the CMG complexes to reconfigure and bypass each other as double-stranded DNA is unwound bi-directionally^5, 6^. As DNA synthesis proceeds, Mcm10 stabilizes the replisome to prevent replication stress and promote genome stability^7–9^.

During oncogenic transformation, cells overexpress replication factors to drive proliferation^10, 11^. Therefore, it is not surprising that *MCM10* is commonly upregulated in cancer cell lines and tumor samples, observations that suggest that these cells rely on Mcm10 to prevent genomic instability from reaching lethal levels^10^. Until recently these observations constituted the only examples of *MCM10*-associated human disease. Recently, we identified human germline *MCM10* mutations in two unrelated families that were associated with distinct phenotypes: natural killer (NK) cell deficiency (NKD)^12^ and restrictive cardiomyopathy (RCM) associated with thymic and splenic hypoplasia. Both of these conditions appear to be the result of compound heterozygous *MCM10* mutations. Previous studies of human *MCM10* have relied heavily on overexpression of epitope-tagged constructs and/or transient knockdown, and thus poorly recapitulate the effects of these patient mutations^10^. To gain a more accurate understanding of the cellular phenotypes underlying *MCM10*-associated NKD or RCM, we chose to model these mutations in human cell lines.

Here, we demonstrate that human *MCM10* is haploinsufficient in transformed HCT116 and non-transformed telomerase immortalized hTERT RPE-1 cells (referred to subsequently as RPE-1). These phenotypes were more severe in HCT116 cells, disrupting normal cell cycle distribution and affecting global DNA synthesis due to decreased origin firing. Chronic Mcm10 deficiency in both cell types caused increased cell death and revealed an unexpected telomere maintenance defect. Our data suggest that telomeric replication fork stalling compromised telomerase access to chromosome ends. Telomere erosion was not a characteristic of *CDC45* or *MCM4* haploinsufficiency, suggesting that the telomeric replisome is uniquely reliant on robust Mcm10 levels. Finally, our data demonstrate that stalled forks required nuclease processing to prevent fork collapse, as *MCM10*:*MUS81* double mutants displayed exacerbated growth and telomere maintenance phenotypes. Taken together, our results revealed that Mcm10 is critical for human telomere replication and suggest that defective telomere maintenance caused both *MCM10-*associated NKD and RCM.

## RESULTS

### Modeling *MCM10* patient mutations reveals haploinsufficiency

Compound heterozygous *MCM10* mutations were identified in unrelated patients that presented with NKD or fetal RCM with thymic and splenic hypoplasia (Table 1). Clinical and genetic analysis of the NKD patient and family was described previously^12^. The NKD-associated mutations were identified in a single patient and included one missense (c.1276C>T, p.R426C) and one nonsense (c.1744C>T, p.R582X) mutation. Clinical and genetic analysis of the RCM patients and family is described in the Supplementary Note. The RCM-associated mutations were identified in three affected siblings and included one splice donor site (c.764+5G>A, p.D198GfsTer10) and one nonsense (c.236delG, p.G79EfsTer6) mutation (Fig. 1a,b, Supplemental fig. 1a,b). Homozygous individuals were not reported on the Genome Aggregation Database (gnomAD)^13^ for any of the patient-associated *MCM10* mutations. Furthermore, allele frequencies reported for these mutations are consistent with very rare, autosomal recessive alleles (Table 1). Both the NKD-associated missense and RCM-associated splice site *MCM10* mutations map to the conserved Mcm10 internal domain (ID; Supplemental fig. 1c). The NKD-associated p.R426C substitution resides C-terminal to zinc-finger 1 and was not predicted to significantly alter protein structure (Figure 1c). Consistent with this, RPE-1 cells homozygous for this mutation showed stable Mcm10 expression, although analyses of proliferation and sensitivity to ultra-violet (UV) light demonstrated that this allele is hypomorphic^12^.

**Fig. 1.**
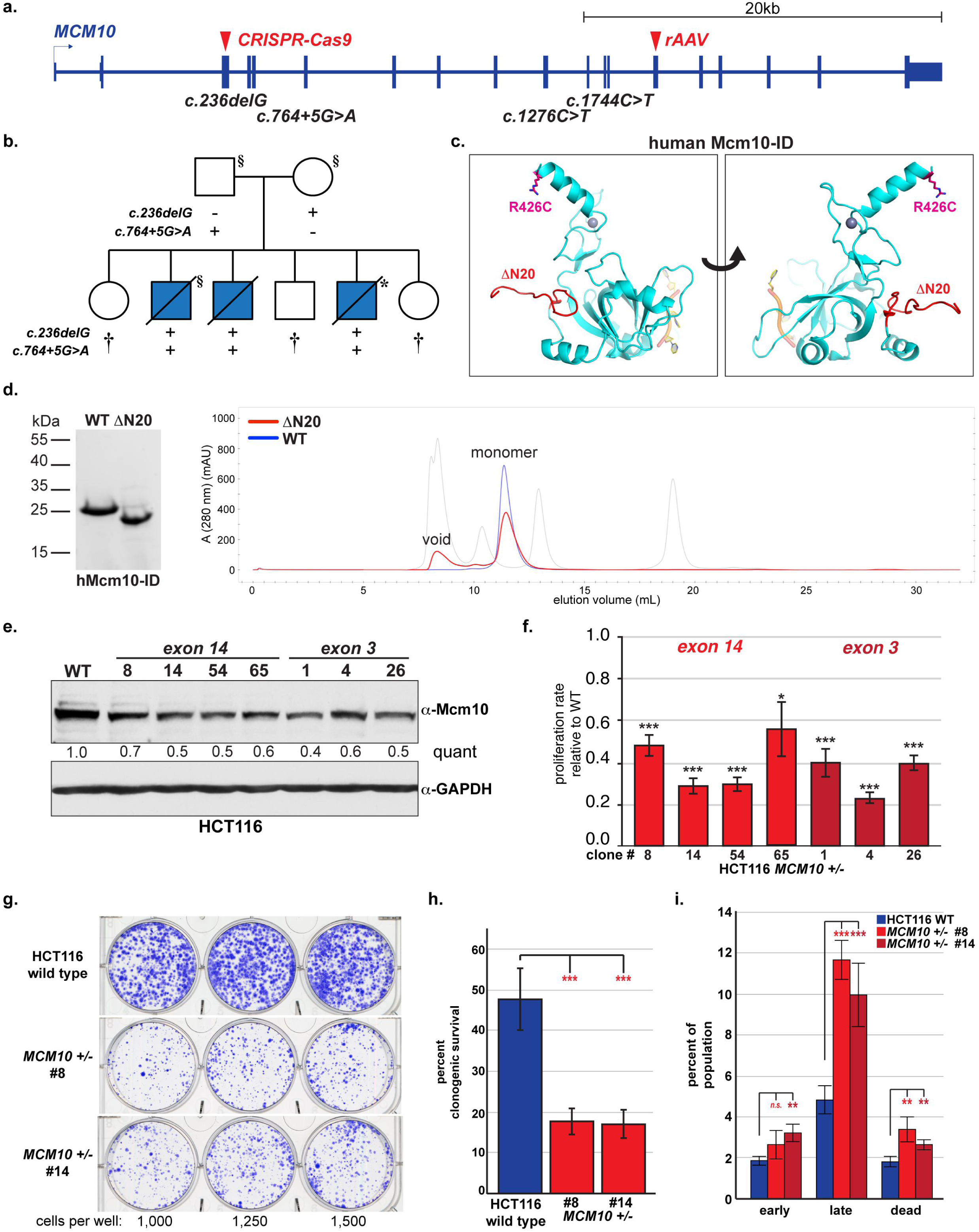
Modeling *MCM10* patient-associated mutations *in vitro* and *in vivo*. **a)** Schematic of human *MCM10* indicating NK- and RCM-associated patient mutations (annotated based on *MCM10* transcript NM_018518.5) and exons targeted using CRISPR-Cas9 (exon 3) or rAAV (exon14) to generate *MCM10^+/-^* cell lines. **b)** Pedigree and segregation of *MCM10* mutations in RCM family. Blue shading indicates fetal RCM. Individuals that underwent exome (§) or genome (*) sequencing are indicated. Clinically unaffected children that are not carriers of both pathogenic mutations (†) are indicated (carrier status of minors thereby not disclosed). **c)** Structural model of the human Mcm10-ID with a bound single-stranded DNA, based on *Xenopus laevis* Mcm10-ID (Protein Data Bank accession codes 3EBE^68^ and 3H15^69^). The zinc ion is shown as a gray sphere. The locations of the R426C and *<ι>Δ</i>*N20 mutations are indicated. **d)** (Left) Coomassie blue-stained SDS-PAGE gel of purified WT and *<ι>Δ</i>*N20 Mcm10-ID. (Right) Size exclusion chromatography profiles comparing elution of WT and *<ι>Δ</i>*N20 Mcm10-ID. The molecular weight standard (gray) included thyroglobulin (670 kDa), *<ι>γ</i>*-globulin (158 kDa), ovalbumin (44 kDa), myoglobin (17 kDa), and vitamin B12 (1.3 kDa). **e)** Western blot for Mcm10 with GAPDH as a loading control. Quantification of Mcm10 levels normalized to loading control, relative to wild type is indicated. **f)** Average proliferation rate in *MCM10^+/-^* cells normalized to wild type. For each cell line n = 6 wells across three biological replicates. **g)** Comparison of clonogenic survival of HCT116 wild type (top) and *MCM10^+/-^* cells (middle/bottom). Cells plated per well are noted. **h)** Percentage clonogenic survival in HCT116 wild type (blue) and *MCM10^+/-^* cells (red), n = 15 wells across ten biological replicates. **i)** Average percentage of each population represented by early apoptotic, late apoptotic or dead cells. HCT116 wild type (blue) and clonal *MCM10^+/-^* cell lines (red) are shown. Error is indicated in **f, h** and **i** as standard deviation and significance was calculated using two-tailed student’s *t-test* with *<.05; **<.01, ***<.001.

**Table 1.**

| **Mutation** | **Pathology** | **Functional Class** | **gnomAD frequency** |
| --- | --- | --- | --- |
| c.1276C>T, p.R426C<br>c.1744C>T, p.R582X | NK cell deficiency | hypomorph<br>LOF | 7/282,582<br>n.r. |
| c.764+5G>A,p.D198GfsTer10<br>c.236delG, p.G79EfsTer6 | Cardiomyopathy with<br>lymphoreticular hypoplasia | hypomorph<br>LOF | 1/232,426<br>n.r. |

Mutations annotated in reference to *MCM10* transcript NM\_018518.5. Frequencies based on gnomADv.2.1.1<sup>13</sup>. LOF = loss of function, n.r. = not reported.

The RCM-associated c.764+5G>A mutation altered a splice donor site that caused exclusion of exon 6 from the mature mRNA and results in expression of Mcm10 carrying a deletion of amino acids 198-255 (Supplemental fig. 1b,c). The first 38 amino acids of this region are part of the unstructured linker domain that connects the N-terminus with the ID. The adjacent 20 amino acids are part of the OB-fold. To gain insight into how this internal deletion affected Mcm10 function, we compared the biochemical behaviors of the hMcm10-ID (residues 236-435) and a corresponding N-terminally truncated mutant encompassing amino acids 256-435 (ΔN20; Fig. 1d). Size-exclusion chromatography showed that, although both proteins were predominantly monomeric, ΔN20 was partially aggregated and eluted in the void volume (Fig 1d). In addition, to assess protein secondary structure we recorded circular dichroism (CD) spectra at 25 °C for the wild type and ΔN20 ID. These spectra showed pronounced negative ellipticity peaks at 205 nm, which is typical for a zinc finger protein^14^ (Supplemental fig. 1d) and suggested that these proteins had similar secondary structures. The spectrum of the wild type protein at 60 °C remained mostly unchanged, indicating a stable protein structure. However, the spectrum of the ΔN20 mutant had a significantly reduced 205 nm peak at 60 °C, indicative of decreased thermal stability (Supplemental fig. 1d). Next, we collected thermal denaturation data by monitoring CD signals at 205 nm in a temperature range of 40 to 90 °C. From the single wavelength curves the melting temperatures (T_m_) of wild type hMcm10-ID and the mutant ΔN20 were determined to be 71 °C and 56 °C, respectively (Supplemental fig. 1e). Overall, these experiments demonstrated that the N-terminal deletion leads to compromised stability of hMcm10-ID and makes the protein prone to aggregation. Taken together, these data suggest that the *MCM10* missense and splice donor mutations associated with NKD and RCM, respectively, result in the expression of abnormal proteins, with the RCM-associated mutation causing a more severe reduction in Mcm10 function.

The hypomorphic mutations caused pathological defects only in combination with an *MCM10* nonsense mutation. To understand the impact of these nonsense mutations on each patient’s disease, we modeled them in HCT116 cells, a cell line amenable to multiple gene targeting strategies. Initially, we utilized rAAV (recombinant adeno-associated virus) to delete exon 14 and introduce a premature stop codon resulting in a p.583X mutation to model the NKD-associated p.582X mutation (Fig. 1a). Whereas CRISPR-Cas9 (clustered regularly interspersed palindromic repeat-CRISPR-associated 9) was used to introduce a premature stop codon in exon 3 to model the RCM-associated mutation in the same exon (Fig. 1a). Due to the introduction of premature stop codons, these mutations presumably led to nonsense-mediated mRNA decay. Regardless, if translated, Mcm10 would be truncated prior to the nuclear localization sequence and thus remain cytoplasmic^15^. Western blot analysis demonstrated stable Mcm10 reduction in *MCM10^+/-^* cell lines (Fig. 1e), which significantly slowed cell proliferation (Fig. 1f) and decreased clonogenic survival (Fig 1g,h) due in part to increased cell death (Fig 1i). These observations demonstrated that in HCT116 cells *MCM10* is genetically haploinsufficient.

### Reduced origin firing causes DNA replication defects and impairs viability of Mcm10-deficient cells

To define the cause of the growth defect in *MCM10^+/-^* mutants, we utilized quantitative analytical flow cytometry. Cell cycle distribution was assessed by staining with DAPI (4’,6-diamidino-2-phenylindole) for DNA content in combination with a pulse of EdU (5-ethynyl-2’-deoxyuridine) to label S-phase cells (Fig. 2a). In *MCM10^+/-^* cells, we detected a significant increase in G1- and decrease in S-phase populations (Fig. 2b) that suggested delayed progression through the G1/S-phase transition. Next, we measured chromatin-bound Mcm2 and EdU-labeled DNA to quantify G1-phase licensing and S-phase DNA synthesis (Fig. 2c-d)^16^. Origin licensing in *MCM10^+/-^* mutants was identical to wild type cells (Fig. 2e). However, we observed a significant DNA synthesis defect during the 30-minute labeling pulse in *MCM10^+/-^* cells (Fig. 2f). These data demonstrated a requirement for Mcm10 in DNA synthesis, but not origin licensing, and are consistent with the published roles for Mcm10 in DNA replication^10^.

**Fig. 2.**
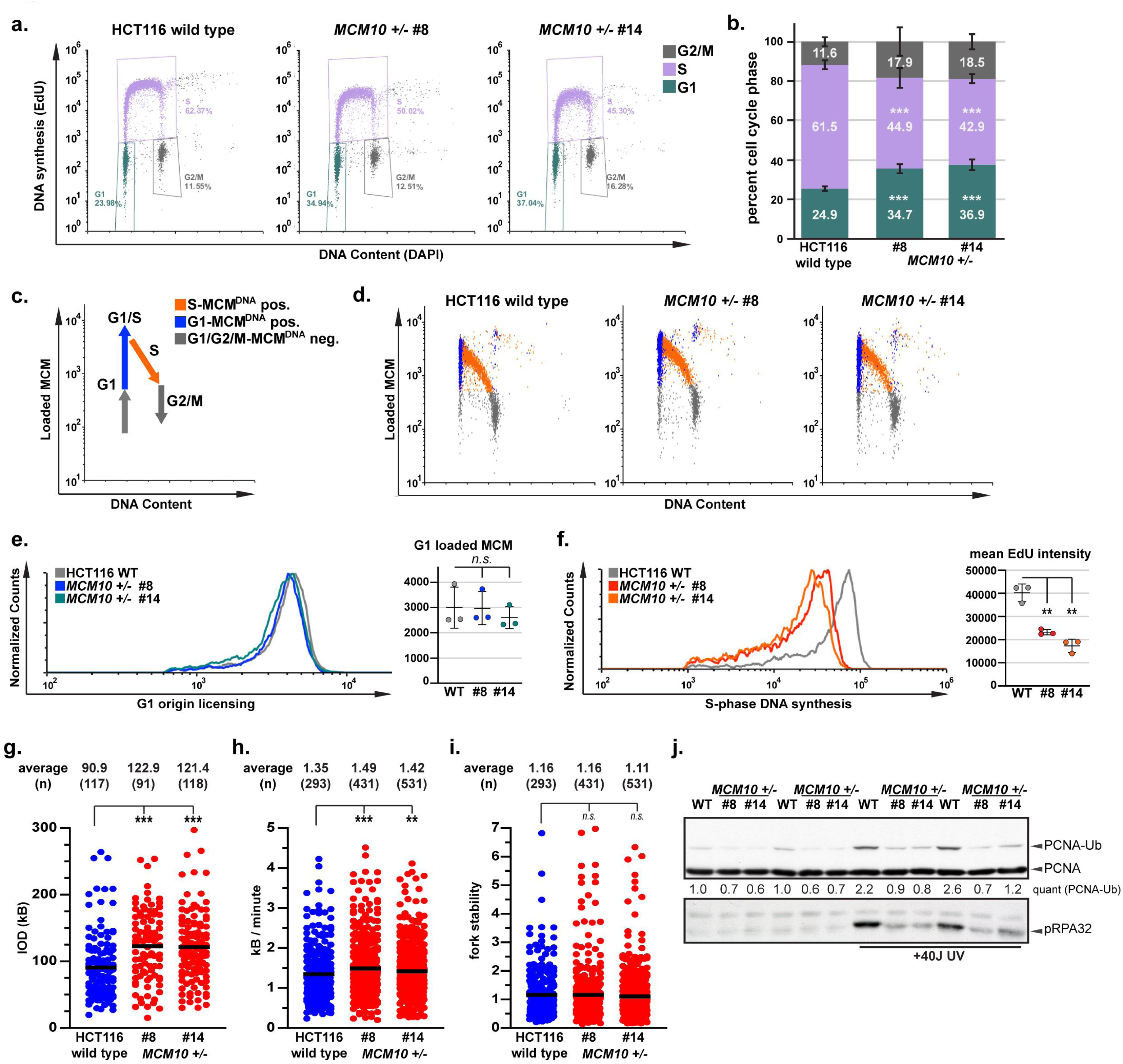
Mcm10 deficiency causes significant cell cycle and DNA synthesis defects. **a)** Cell cycle distribution of HCT116 wild type and *MCM10^+/-^* cells. Percentage of each population in G1-(green), S-(purple) or G2/M-phase (gray) is shown. **b)** Cell cycle distribution of HCT116 wild type and *MCM10^+/-^* cell lines from three biological replicates. Percentage of each population in G1-(green), S-(purple) and G2/M-phase (gray) is shown. **c)** Schematic of flow cytometry analysis. Cell-cycle phase is defined by DNA content, EdU incorporation and chromatin loaded MCM2. **d)** Analytical flow cytometry plots for HCT116 wild type and *MCM10^+/-^* cells. G1-phase/MCM positive cells (blue), S-phase/MCM positive cells (orange) and G1- or G2/M-phase/MCM negative cells (gray) are indicated. **e)** Comparison of origin licensing (left) and quantification of G1 loaded MCM2 (n = 3) in HCT116 wild type (gray) and *MCM10^+/-^* cells (blue/green). **f)** Comparison of S-phase DNA synthesis (left) and mean EdU intensity (n = 3) in HCT116 wild type (gray) and *MCM10^+/-^* cells (red/orange). Error bars in **b, e** and **f** indicate standard deviation and significance was calculated using two-tailed student’s *t-test* with *<.05; **<.01, ***<.001. **g)** Inter-origin distance (IOD) quantification from three technical replicates across two biological replicates in wild type (blue) and *MCM10^+/-^* cells (red). Average IOD and number (n) quantified is listed. **h)** Fork speed from three technical replicates across two biological replicates in wild type (blue) and *MCM10^+/-^* cells (red). Average fork speed (kb/minute) and number (n) quantified is listed. **i)** Fork stability from three technical replicates across two biological replicates in wild type (blue) and *MCM10^+/-^* cells (red). Average fork stability and number (n) quantified is listed. Statistical significance for **g-i** was calculated using Mann-Whitney Ranked Sum Test with *<.05; **<.01, ***<.001. **j)** Chromatin associated PCNA, PCNA-Ub and phospho-RPA32, which binds to ssDNA exposed during replication stress, with and without 40 J UV treatment. Quantification of PCNA-Ub levels normalized to unmodified PCNA, relative to the first lane wild type sample is indicated.

To understand the DNA synthesis defect in *MCM10^+/-^* cells, we performed DNA combing. We first measured inter-origin distance (IOD) to determine if origin firing was perturbed. Whereas wild type cells displayed an average IOD of ∼91 kb, the IOD in *MCM10^+/-^* cells markedly increased to ∼120 kb (Fig. 2g). This difference equates to ∼25% fewer origin firing events in *MCM10* mutants. Next, we measured global fork speed and stability. Fork speed was modestly, but significantly, increased in *MCM10^+/-^* cells (Fig. 2h), consistent with the inverse regulation of fork speed and origin firing in eukaryotes^17, 18^. Fork stability was not changed (Fig. 2i). These data suggested that Mcm10 deficiency reduced the number of active forks. To confirm this idea, we assessed ubiquitination (Ub) of proliferating cell nuclear antigen (PCNA), because this modification at lysine 164 occurs specifically at active forks in response to replication stress^19^. Since HCT116 cells exhibit intrinsic replication stress, PCNA-Ub is detectable under unperturbed conditions (Fig. 2j)^20^. To enhance PCNA-Ub, we exposed cells to 40 J/m^2^ of UV radiation. As expected, PCNA-Ub was 1.5-to 3-fold higher in wild type cells than in *MCM10^+/-^* cells (Fig. 2j). Furthermore, phosphorylated RPA32 was elevated in wild type cells in comparison to *MCM10^+/-^* cells exposed to UV radiation. Therefore, Mcm10 deficiency decreased origin firing and resulted in fewer active forks. However, forks were stable globally and did not elicit an increased stress response.

### Replication stress in Mcm10-deficient cells causes genomic instability and spontaneous reversion of the *MCM10* mutation

To understand if chronic Mcm10 deficiency promoted genome instability, we generated late passage HCT116 wild type and *MCM10^+/-^* populations. As *MCM10^+/-^* cells were passaged, we observed an increase in cells with abnormal morphology and multiple enlarged or pyknotic nuclei indicative of decreased viability (Supplementary Fig. 2a). Karyotype analysis of wild type cells after ∼200 population doublings (PDs) as well as *MCM10^+/-^* populations after mid-(∼25 PDs) and late passage (∼100 PDs) revealed increased genome instability in the mutants (Supplementary Fig. 2b). Late passage wild type cells harbored three populations represented by the parental HCT116 karyotype and two derivatives, with 10% of karyotypes carrying unique aberrations (Supplementary table 1). Mid-passage *MCM10^+/-^* cells were comprised of one major population distinct from the parental karyotype carrying two novel aberrations, as well as seven additional novel karyotypes (Supplementary table 1). Overall, 23% of mid-passage *MCM10^+/-^* karyotypes had unique aberrations. Furthermore, late passage *MCM10^+/-^* cells consisted of a clonal population distinct from the parental karyotype carrying three novel aberrations, as well as nineteen additional novel karyotypes (Supplementary table 1). Overall, 63% of late-passage *MCM10^+/-^* karyotypes had unique aberrations, suggesting that they were the result of independent translocations. The acquired chromosomal rearrangements in *MCM10* mutants significantly overlapped with common fragile sites (CFSs) (80%; Supplementary table 1)^21^, which is consistent with a role for Mcm10 in preventing fragile site breakage^22^.

The overlap of translocation hot spots with CFSs prompted us to investigate telomere maintenance, as telomeres are also origin-poor and hard-to-replicate regions^23^. We performed telomere restriction fragment (TRF) length analysis to measure average length over time. Whereas wild type telomeres were stable, telomeres in *MCM10^+/-^* cell lines were shorter at early passage and eroded over time (Fig. 3a). Consistent with these data, early passage exon 3 *MCM10^+/-^* HCT116 cells contained eroded telomeres (Supplementary Fig. 2c). To independently confirm this phenotype, we performed telomere fluorescence *in situ* hybridization (t-FISH) of metaphase chromosomes. Consistent with our TRF analyses we observed a significant increase in chromosomes lacking a telomere signal (“signal free ends”) in *MCM10^+/-^* metaphase spreads (Fig 3b). We also quantified fragile telomeres, chromosome ends with multiple telomere foci that are indicative of abnormal structure^24^, but did not find any increase in *MCM10^+/-^* mutants (Fig. 3b). Next, we measured *<ι>β</i>*-galactosidase (*<ι>β</i>*-gal) activity to determine whether *MCM10^+/-^* cells activated cellular senescence pathways^25, 26^. We observed significantly higher activity in mutant cell extracts, further corroborating the telomere maintenance defect (Supplementary Fig. 2d). Finally, we measured telomerase activity using the telomeric repeat amplification protocol (TRAP)^27^, which ensured that activity was equivalent in all cell lines (Fig. 3c). We could thus exclude telomerase inactivation as the cause of telomere erosion.

**Fig. 3.**
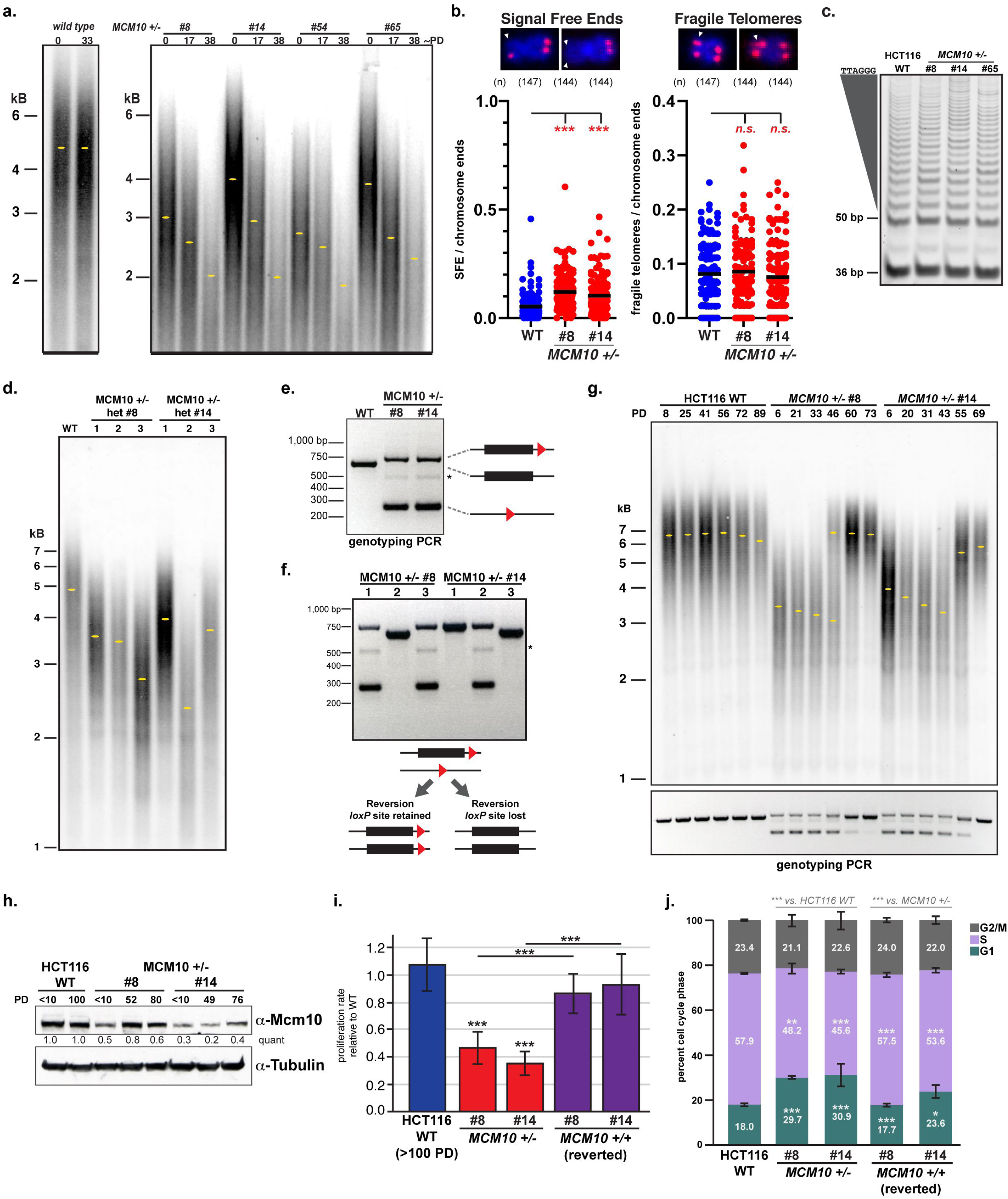
Mcm10 deficiency causes telomere erosion that is rescued following spontaneous reversion of the *MCM10* mutant locus. **a)** TRF analysis of HCT116 wild type (left) and *MCM10^+/-^* cells (right). Estimated PDs is indicated. Yellow dots indicate the location of peak intensity. **b)** Signal free-ends (left) and fragile telomeres (right) in HCT116 wild type (blue) and *MCM10^+/-^* cells (red). Statistical significance was calculated using two-tailed student’s *t-test* with ***<.001; n = metaphases analyzed per cell line. Scale bars are 1 µm. **c)** TRAP assay from HCT116 wild type (left) and *MCM10^+/-^* cells (right). The internal PCR control at 36 bp and telomerase above 50 bp are noted. **d)** TRF analysis of HCT116 wild type (left) and independent late passage *MCM10^+/-^* cell populations from two parental *MCM10^+/-^* cell lines (right). Yellow dots indicate the location of peak intensity. **e)** *MCM10^+/-^* exon 14 genotyping of alleles carrying a *loxP* site 3’ of exon 14 (upper band) or a *loxP* scar (lower band), in comparison to the wild type locus (middle band). A faint non-specific band is noted (asterisk). **f)** *MCM10^+/-^* exon 14 genotyping PCR in late passage *MCM10^+/-^* populations showing alleles that have one *loxP* site 3’ of exon 14 (upper band) or a *loxP* scar (lower band), as well as exon 14 reverted alleles that have retained or lost the 3’ *loxP* site. A faint non-specific band can also be detected (asterisk). **g)** TRF analysis in HCT116 wild type and *MCM10^+/-^* cells (top). PDs for each population are shown. Yellow dots indicate the location of peak intensity. Genotyping PCR for each TRF sample is shown (bottom). **h)** Western blot analyses for Mcm10 with tubulin as a loading control. Quantification of Mcm10 levels normalized to loading control, relative to the first lane wild type sample is indicated. PDs for each cell line are noted. **i)** Proliferation rate in HCT116 wild type, *MCM10^+/-^* and reverted cells normalized to early passage wild type cells, for each cell line n = 6 replicate wells across two biological replicates. **j)** Cell cycle distribution of HCT116 wild type, *MCM10^+/-^* and reverted cell lines, n = 4 across two biological replicates. Percentage of each population in G1-(green), S-(purple) and G2/M-phase (gray) is shown. Error bars in **h** and **i** indicate standard deviation and significance was calculated using two-tailed student’s *t-test* with *<.05; **<.01, ***<.001.

To understand whether Mcm10 deficiency might drive cells into telomere crisis, we propagated six independent *MCM10^+/-^* populations for >75 PDs. TRF analyses documented that these populations contained telomeres 2 to 4 kb in length (Fig. 3d). Consistent with these data, the frequency of signal free ends remained elevated in *MCM10^+/-^* cells, but was not increased in comparison to early passage mutants (Supplementary Fig. 2g, Fig. 3b). Again, we did not detect changes in telomere fragility (Supplementary Fig. 2g). These observations suggested that short telomere length stabilized in late passage *MCM10^+/-^* populations. To evaluate this idea, we first confirmed the genotype of each population. Unexpectedly, only three populations carried the heterozygous PCR pattern and three populations had spontaneously reverted the exon 14 mutations (Fig. 3e,f). Reversion resulted in two alleles carrying wild type coding sequence. Some cell lines retained the 3’ *loxP* site on both alleles, while others lost the *loxP* sites completely (Fig. 3e,f). These results prompted us to conduct additional long-term experiments closely monitoring PDs, telomere length and genotype. Analysis of these time courses revealed two novel spontaneous reversion events (Fig. 3g). Importantly, *MCM10* reversion corresponded with rescued telomere length (Fig. 3g), increased Mcm10 levels (Fig. 3h), growth rate recovery (Fig. 3i) and rescued defects in cell cycle distribution (Fig. 3j).

To confirm that reversion events were not the misinterpretation of culture contamination with wild type cells, we marked five independent *MCM10^+/-^* populations with a puromycin (PURO) resistance gene. Following drug selection, the genotype of each population was confirmed as heterozygous and cell lines were propagated and collected at regular time intervals. Each population underwent additional PURO selection twice during the time course (Supplementary Fig. 2h). After ∼100 PDs, one of the five populations spontaneously reverted (Supplementary Fig. 2i). In this population, telomeres eroded initially but recovered and stabilized following reversion. Taken together, our data demonstrate that the toxic nature of Mcm10 deficiency was actively selected against, as *MCM10^+/-^* cells spontaneously reverted the locus to restore the *MCM10* coding sequence and rescue mutant phenotypes.

### Deficiency of essential replisome proteins Cdc45 or Mcm4 does not cause telomere erosion

The mutant phenotypes in *MCM10^+/-^* cells led us to ask if HCT116 cells are broadly sensitive to heterozygosity of essential replisome genes. To test this, we constructed *CDC45^+/-^* and *MCM4^+/-^* cell lines. These genes encode CMG helicase proteins and contribute to the same processes that require Mcm10^10^. *CDC45^+/-^* and *MCM4^+/-^* cell lines showed reduction in the protein corresponding to the inactivated gene, but no change in Mcm10 levels (Supplementary Fig. 3a). Unexpectedly, Cdc45 levels were significantly reduced in *MCM4^+/-^* mutants, suggesting that Mcm4 stabilizes Cdc45 protein. We also confirmed that Cdc45 and Mcm4 levels remained normal in *MCM10^+/-^* cell lines (Supplementary Fig. 3b). *CDC45^+/-^* and *MCM4^+/-^* populations proliferated slower than wild type cells (Supplementary Fig. 3c), although this phenotype was not as severe as in *MCM10^+/-^* cells (Fig. 1d). The amount of chromatin-bound PCNA-Ub was similar in *CDC45^+/-^* and *MCM4^+/-^* mutants in comparison to wild type cells, implying that the number of active forks was not reduced, unlike our observations in *MCM10^+/-^* cells (Supplementary Fig. 3d, Fig. 3e). Finally, TRF analyses revealed that telomere length was stable in *CDC45^+/-^* and *MCM4^+/-^* mutants (Supplementary Fig. 3e-f). Taken together, these data argue that HCT116 cells are sensitive to inactivation of one *CDC45* or *MCM4* allele, but they are significantly more sensitive to *MCM10* heterozygosity that is associated with a unique telomere maintenance defect.

### Modeling *MCM10* patient mutations in non-transformed RPE-1 cells confirms defects in telomere maintenance

*MCM10* haploinsufficiency was unexpected, as heterozygous mice were healthy and fertile^28^ and haploinsufficiency is uncommon among human genes^29^. Moreover, the requirement for bi-allelic *MCM10* mutation to elicit a clinical phenotype suggested that haploinsufficiency might be unique to cancer cells. To evaluate this idea, we modeled the *MCM10* exon 3 patient nonsense mutation (c.236delG, p.G79EfsTer6) in RPE-1 cells. *MCM10^+/-^* RPE-1 cells showed stable reduction of Mcm10 (Fig. 4a), which significantly slowed cell proliferation (Fig. 4b) and increased cell death (Fig 4c). Unexpectedly, significant changes in the cell cycle distribution of *MCM10^+/-^* cells were not observed (Fig. 4e-f) and the amount of origin licensing (Fig. 4g-h) and S-phase DNA synthesis (Fig. 4g,i) were normal. Notably, origin licensing and DNA synthesis were significantly higher in wild type HCT116 than in wild type RPE-1 cells (Fig. 4i-j). Presumably, these data reflect differences between cell types, including changes that HCT116 cells underwent during oncogenic transformation, and suggest that highly proliferative transformed cells may be inherently more sensitive to Mcm10 deficiency.

**Figure 4.**
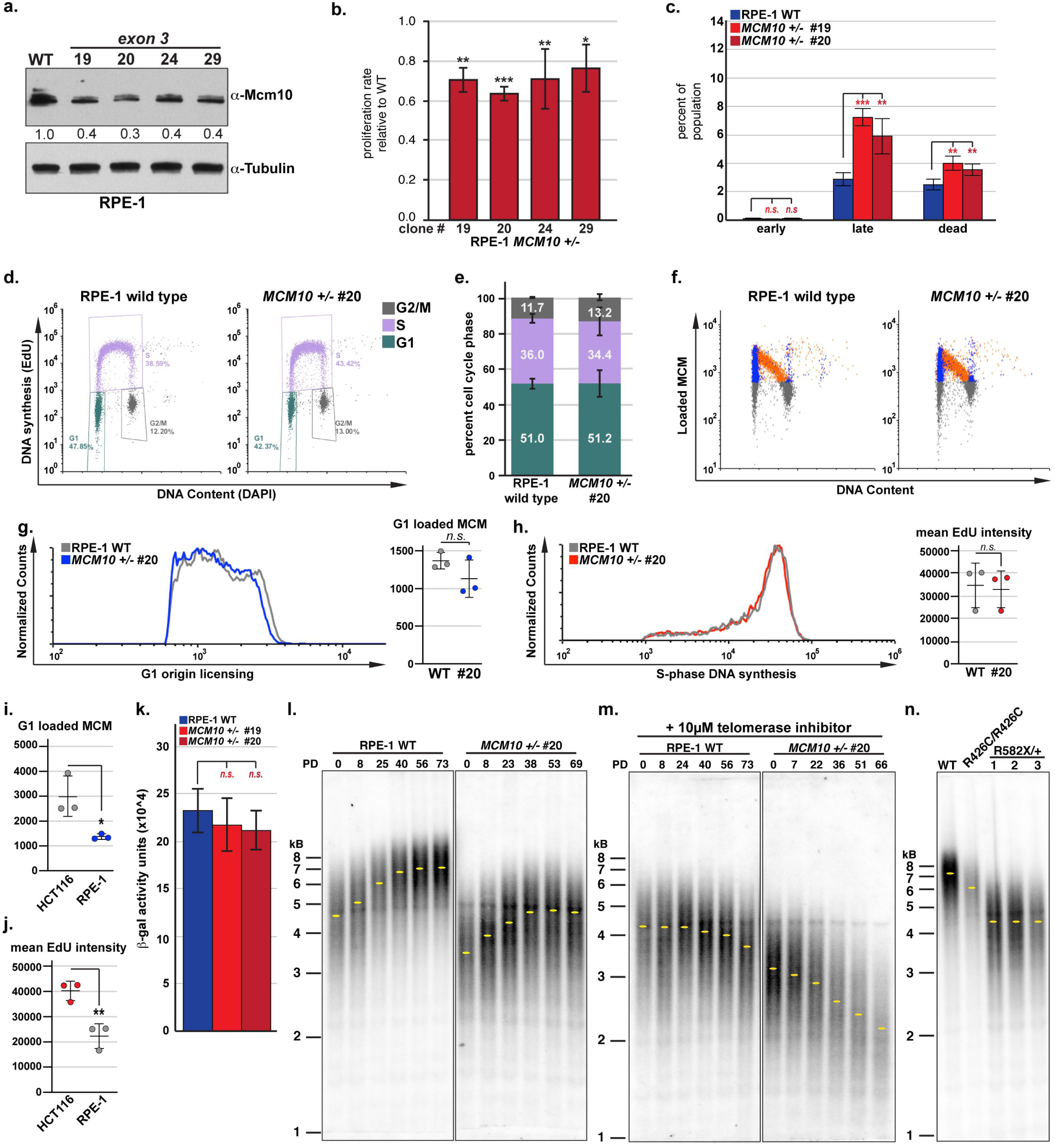
*MCM10* heterozygous RPE-1 cell lines have reduced Mcm10 expression, impaired proliferation and telomere maintenance defects. **a)** Western blot for Mcm10 with tubulin as a loading control. Quantification of Mcm10 levels normalized to loading control, relative to wild type is indicated. **b)** Average proliferation rate in *MCM10^+/-^* cells normalized to wild type. For each cell line n = 6 wells across three biological replicates. **c)** Average percentage of each population represented by early apoptotic, late apoptotic or dead cells. RPE-1 wild type (blue) and clonal *MCM10^+/-^* cell lines (red) are shown. Error bars indicate standard deviation and statistical significance was calculated using student’s *t-test* with *<.05; **<.01, ***<.001; n = 4 for all data points. **d)** Cell cycle distribution of RPE-1 wild type and *MCM10^+/-^* cells. Percentage of each population in G1-(green), S-(purple) and G2/M-phase (gray) is shown. **e)** Cell cycle distribution of RPE-1 wild type and *MCM10^+/-^* cells from three biological replicates. Percentage of each population in G1-(green), S-(purple) and G2/M-phase (gray) is shown. **f)** Flow cytometry plots for RPE-1 wild type and *MCM10^+/-^* cells. G1-phase/MCM positive cells (blue), S-phase/MCM positive cells (orange) and G1- or G2/M-phase/MCM negative cells (gray) are indicated. **g)** Comparison of origin licensing (left) and G1 loaded MCM2 (n = 3) in RPE-1 wild type (gray) and *MCM10^+/-^* cells (blue). **h)** Comparison of S-phase DNA synthesis (left) and mean EdU intensity (n = 3) in RPE-1 wild type (gray) and *MCM10^+/-^* cell lines (red). **i)** G1 loaded Mcm2 (n = 3) in wild type HCT116 (gray) and RPE-1 (blue) cells. **j)** Mean EdU intensity (n = 3) in wild type HCT116 (red) and RPE-1 (gray) cells. **k)** Quantification of *<ι>β</i>*-gal activity expressed as arbitrary fluorescence units normalized to total protein for RPE-1 wild type (blue) and clonal *MCM10^+/-^* cell lines (red). Error bars in **b, c, e,** and **g-k** indicate standard deviation and significance was calculated using two-tailed student’s *t-test* with *<.05; **<.01, ***<.001. **l)** TRF analysis in RPE-1 wild type and *MCM10^+/-^* clone #20. PDs for each cell line are noted. Yellow dots indicate the location of peak intensity. **m)** TRF analysis RPE-1 wild type and *MCM10^+/-^* clone #20 in the presence of telomerase inhibitor. PDs for each cell line are noted. Yellow dots indicate the location of peak intensity. **n)** TRF analysis RPE-1 wild type and *MCM10* mutant cell lines carrying NK-associated patient mutations. Yellow dots indicate the location of peak intensity.

We hypothesized that the slowed proliferation and increased cell death in *MCM10* mutant RPE-1 cells was due to defects in telomere maintenance, as global changes in DNA synthesis were not observed (Fig. 4h). First, we measured *<ι>β</i>*-gal activity to determine whether *MCM10^+/-^* cells activated senescence pathways; however, no increase was detected in *MCM10* mutants (Fig. 4k). When we monitored telomere length in wild type and *MCM10^+/-^* cells at regular intervals we found that telomeres in *MCM10^+/-^* cells were shorter than wild type (Fig. 4l). Surprisingly, we found that telomeres in both cell lines elongated over time (Fig. 4l). This phenotype was telomerase-dependent, as passaging cells with the telomerase inhibitor BIBR1532^30^ prevented elongation (Fig. 4m). These data suggested that telomerase activity in RPE-1 cells was robust enough to catalyze telomere extension, regardless of *MCM10* status. Moreover, although telomeres in wild type RPE-1 cells were stable in the presence of inhibitor, the identical treatment caused rapid erosion in *MCM10^+/-^* mutants (Fig. 4n). These data suggested that patient-associated *MCM10* mutations limited telomere length in RPE-1 cells. To confirm this, we measured average telomere length in RPE-1 cells that carry the homozygous missense (c.1276C>T, p.R426C) or heterozygous nonsense (c.1744C>T, p.R582X) NKD-associated mutations, respectively. Remarkably, each *MCM10* mutant cell line carried telomeres that were significantly shorter than wild type RPE-1 at similar passage (Fig 4n), and the extent of telomere erosion in *MCM10* mutant cells corresponded with the severity of each patient-related mutation. Taken together, our data is consistent with the notion that Mcm10 deficiency limited telomerase-dependent telomere elongation in RPE-1 cells.

### Elevated telomerase activity rescues telomere length but not the inherent replication defect in heterozygous *MCM10* mutant cells

To further evaluate the relationship between telomere length, Mcm10 deficiency and telomerase activity, we cultured wild type and *MCM10* mutant HCT116 cells in the presence of telomerase inhibitor. TRF analyses showed that telomere erosion was exacerbated by telomerase inhibition (Fig. 5a), suggesting that Mcm10 deficiency limited, but did not abolish telomerase-dependent elongation. Interestingly, the population of *MCM10^+/-^* clone #8 was nearly 100% genetically reverted at PD 70, but telomere length had not recovered (Fig. 5a). We continued to propagate this population with and without telomerase inhibitor. Without inhibitor, telomeres efficiently lengthened over time (Fig. 5b). However, with inhibitor present telomeres remained short (Fig. 5b), confirming that telomerase activity was essential for telomere length recovery in reverted *MCM10* cells.

**Fig. 5.**
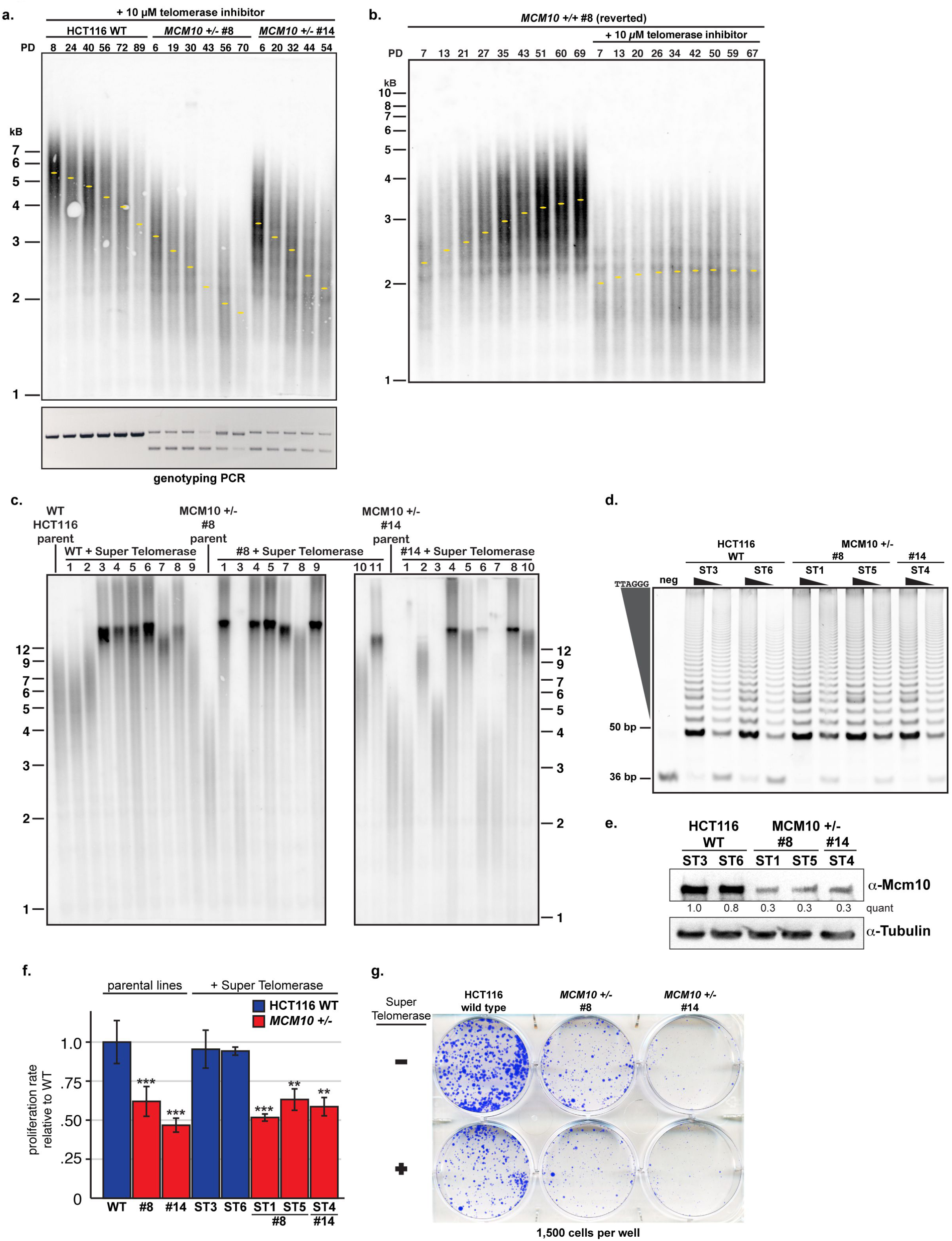
Overexpression of telomerase lengthens telomeres but does not rescue growth and viability defects. **a)** TRF analysis in HCT116 wild type and *MCM10^+/-^* cells (top) in the presence of 10 µM telomerase inhibitor BIBR1532. PDs for each population are shown. Yellow dots indicate the location of peak intensity. Genotyping PCR for each time point is shown (bottom). **b)** TRF analyses in *MCM10^+/-^* clone #8 reverted cells in the presence or absence of telomerase inhibitor. PDs for each population are indicated. Yellow dots indicate the location of peak intensity. **c)** TRF analysis in HCT116 wild type, *MCM10^+/-^* cell lines and ST derivatives of each parental cell line. **d)** Representative TRAP assay comparing telomerase activity in whole cell extracts from HCT116 wild type (left) and *MCM10^+/-^* ST cell lines (right). The internal PCR control at 36 bp and telomerase products beginning at 50 bp are noted. For each cell line, two concentrations of cell extract were used representing a 10-fold dilution. **e)** Western blot analyses of whole cell extracts from HCT116 ST wild type and *MCM10^+/-^* cell lines for Mcm10 with tubulin as a loading control. Quantification of Mcm10 levels normalized to loading control, relative to the first lane wild-type ST3 sample is indicated. **f)** Average proliferation rate in HCT116 wild type, *MCM10^+/-^* and ST cell lines normalized to HCT116 wild type cells. For each cell line n = 4 replicate wells across two biological replicates; error bars indicate standard deviation and statistical significance was calculated using two-tailed student’s *t-test* with *<.05; **<.01, ***<.001. **g)** Comparison of clonogenic survival in HCT116 wild type, *MCM10^+/-^* and ST cell lines. The number of cells plated per well is indicated.

To test whether stable telomerase overexpression could rescue telomere erosion and alleviate HCT116 *MCM10^+/-^* mutant phenotypes, we generated stable cell lines that overexpressed *hTERT* and *hTR* – so-called ‘super-telomerase’ (ST)^31^. Most ST cell lines showed significantly longer telomeres (>12 kb) than observed in normal HCT116 cells (∼4 to 6 kb; Fig. 5c). ST cell lines with significantly extended telomeres and similar telomerase activities (Fig. 5d) were utilized for further experiments. ST expression in *MCM10^+/-^* cell lines did not increase Mcm10 levels (Fig. 5e) nor rescue the diminished proliferation rate (Fig. 5f). In fact, the excessive telomere length slightly decreased clonogenic survival (Fig. 5g). Overall, these data demonstrated that ST expression rescued telomere length without promoting proliferation or viability.

The ST cells allowed us to perform telomeric replication assays similar to those used in models with long telomeres^24^. We utilized DNA combing to specifically analyze telomere and sub-telomere replication (Fig. 6a). In comparison to wild type ST cells, *MCM10^+/-^* ST mutants showed an increase in unreplicated telomeres (Fig. 6b). Moreover, telomeres in *MCM10^+/-^* ST mutants were more often partially replicated, which is consistent with increased fork stalling (Fig. 6c). Next, we used 2-dimensional (2D) gel analyses to detect DNA intermediates associated with telomeric replication stress. A low intensity t-circle arc was observed in all ST cell lines (Fig. 6d). Strikingly, however, *MCM10^+/-^* ST cells significantly accumulated t-complex DNA, which is comprised of branched DNA structures containing internal single-stranded (ss) DNA gaps (Fig. 6d)^32^. Treatment of wild type HCT116 ST cells with hydroxyurea generated t-complex DNA (Fig. 6e), suggesting that these structures are products of replication stress. To further evaluate the nature of t-complex DNA, samples were treated with S1-nuclease to degrade ssDNA. Wild type ST samples were unchanged after S1-nuclease digestion, whereas a significant reduction in t-complex signal occurred in S1-digested *MCM10^+/-^* ST samples (Fig. 6f). These data argued that the accumulated t-complexes in *MCM10* mutants were enriched for ssDNA gaps. Furthermore, the severity of chronic Mcm10 deficiency in ST cells also stimulated spontaneous reversion that rescued the accumulation of t-complex DNA and the proliferation defect (Supplementary Fig. 4a-b). The generation of t-complexes is poorly understood, but their initial characterization was consistent with regressed forks and/or recombination intermediates^32^. To delineate between these possibilities, we measured the frequency of telomere sister chromatid exchanges (t-SCEs) that are produced by homologous recombination. *MCM10^+/-^* ST cells showed a similar frequency of t-SCEs as wild type ST cells (Fig. 6g). These data suggested that t-complexes in Mcm10-deficient cells were not generated by homologous recombination, but are the product of stalled telomeric replication forks.

**Fig. 6.**
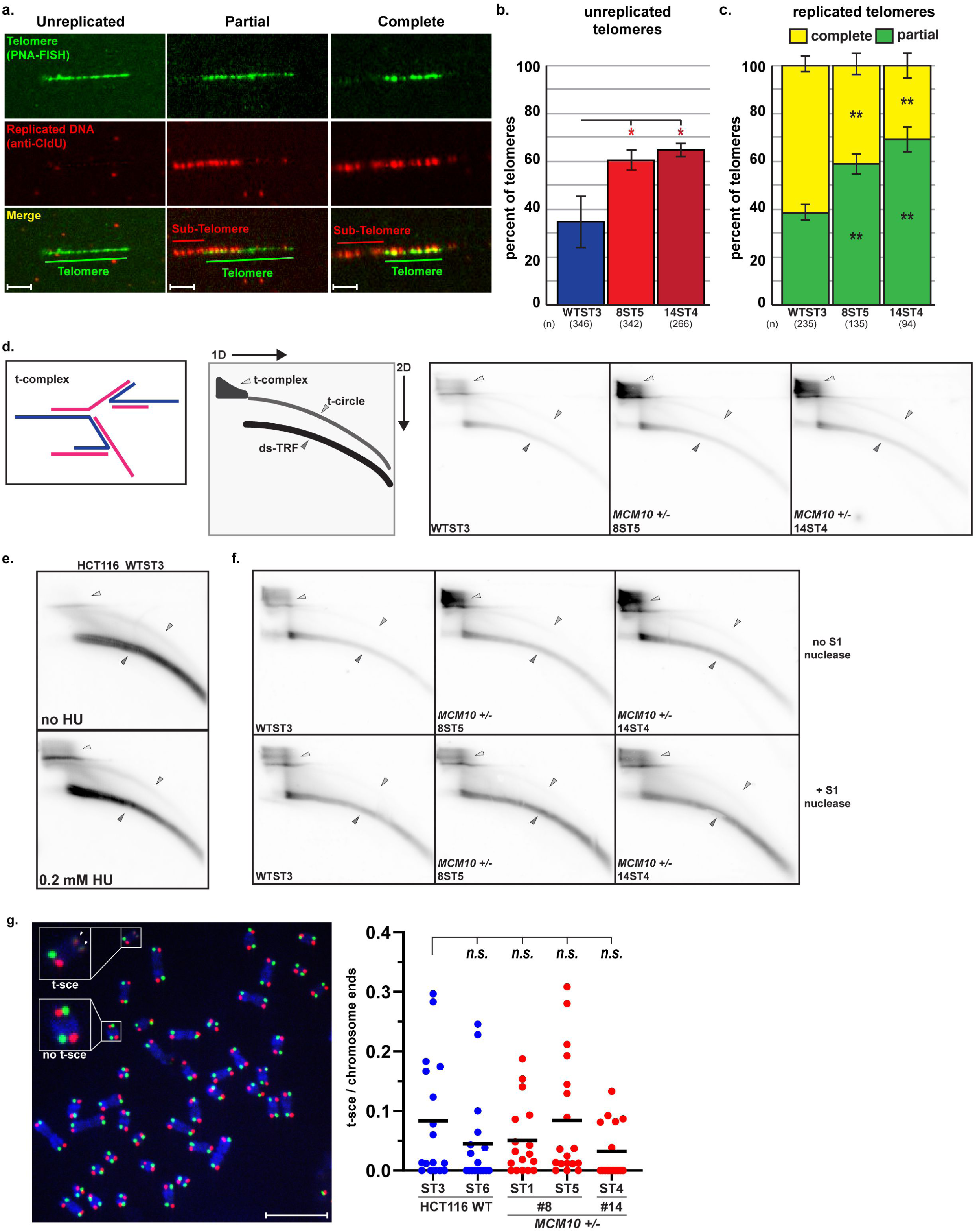
Telomeric replication stress is increased in ST *MCM10^+/-^* HCT116 cells. **a)** Telomere combing images with telomeric DNA (green), nascent DNA (red) and merged images with examples of unreplicated telomeres (left), partially replicated (middle) and completely replicated telomeres (right). Telomeric and sub-telomeric regions are indicated. Scale bars are 3 µm. **b)** Average percentage of unreplicated telomeres in wild type (blue) and *MCM10^+/-^* cell lines (red) ST cell lines, n= total number of telomeres quantified including two or more replicate experiments. **c)** Average percentage of completely replicated (yellow) versus partially/stalled telomeres (green) in ST cell lines, n= total number of replicated telomeres quantified including two or more replicate experiments. Error bars in **b** and **c** indicate standard deviation and significance was calculated using two-tailed student’s *t-test* with *<.05; **<.01, ***<.001. **d)** (Left) Cartoon of t-complex DNA as formerly depicted^32^. (Middle) Diagram of double-stranded telomere restriction fragment (ds-TRF), telomere circle (t-circle) and telomere complex (t-complex) DNA species from 2D gel electrophoresis. (Right) Comparison of 2D gels from ST cell lines. **e)** Comparison of DNA species from 2D gel electrophoresis in HCT116 wild type ST cells with (bottom) and without (top) 4-day HU treatment. **f)** Comparison of DNA species from 2D gels in HCT116 wild type and *MCM10^+/-^* ST cell lines with (bottom) and without (top) S1 nuclease digestion. **g)** (Left) Image of t-SCE staining in ST cell lines. Examples of a t-SCE event and chromosomes without t-SCE are highlighted. (Right) Percentage t-SCE per chromosome ends in ST cell lines. Bars represent median percentage t-SCE per chromosome ends; n = >14 metaphases for each cell line. Significance was calculated using two-tailed student’s *t-test* with *<.05; **<.01, ***<.001. Scale bar is 10 µm.

### Loss of Mus81 exacerbates viability and telomere maintenance defects in Mcm10-deficient cells

One mechanism to resolve fork stalling is the recruitment of structure-specific endonucleases (SSEs) to cleave replication intermediates and stimulate restart. A major player in this pathway is the Mus81 (mutagen-sensitive 81) endonuclease, which functions in complex with Eme1 or Eme2 (essential meiotic SSE1 or 2), as well as a larger DNA-repair tri-nuclease complex^33^. We hypothesized that *MCM10* mutants rely on SSE-dependent fork cleavage to overcome fork stalling. To test this, we generated *MCM10^+/-^*:*MUS81^-/-^* double mutants. Wild type and *MCM10^+/-^* cells expressed equivalent Mus81 protein levels, whereas Mus81 was not detectable in *MUS81^-/-^* mutants (Fig. 7a). Importantly, Mcm10 expression was not altered following the knockout of *MUS81* (Fig. 7a). *MUS81^-/-^* cells showed a growth defect, although not as severe as seen in *MCM10^+/-^* mutants (Fig. 7b). Furthermore, this defect was significantly worse in the double mutants, which proliferated ∼2-fold slower than *MCM10^+/-^* single mutants (Fig. 7b). Comparison of clonogenic survival yielded similar results, wherein *MUS81^-/-^* cells showed a colony formation defect that was less severe than *MCM10^+/-^* mutants and *MCM10^+/-^*:*MUS81^-/-^* cells showed a stronger phenotype than either single mutant (Fig. 7c). Increased apoptosis or cell death was not detected in *MUS81* single mutants (Fig. 7d). However, we measured a significant increase in double mutant cell death (Fig. 7d).

**Fig. 7.**
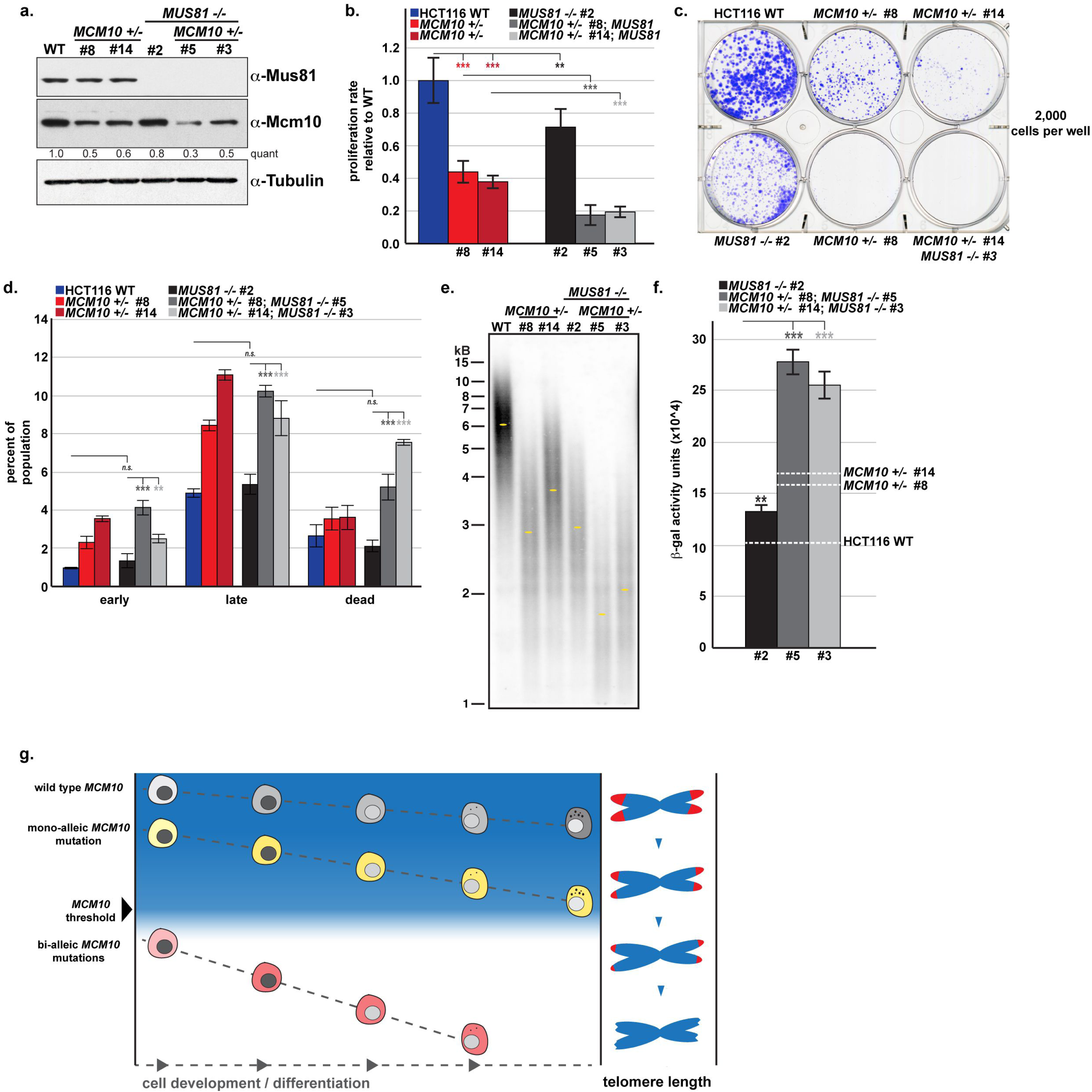
Loss of Mus81 increases the severity of proliferation, viability and telomere length defects in *MCM10^+/-^* cell lines. **a)** Western blot analyses for Mus81 and Mcm10 with GAPDH (left) as a loading control. Quantification of Mcm10 levels normalized to tubulin loading control, relative to the first lane wild type sample is indicated. **b)** Proliferation rate in HCT116 wild type, *MUS81^-/-^*, *MCM10^+/-^* and double mutant cell lines normalized to HCT116 wild type, n = 6 replicate wells across three biological replicates. **c)** Comparison of clonogenic survival of HCT116 wild type, *MUS81^-/-^*, *MCM10^+/-^* and double mutant cell lines. **d)** Average percentage of each population represented by early apoptotic, late apoptotic or dead cells in HCT116 wild-type, *MUS81^-/-^*, *MCM10^+/-^* and double mutant cell lines, n = 2 replicates for HCT116 wild type and *MCM10^+/-^* single mutants; n = 4 replicates across two biological replicates for all *MUS81^-/-^* cell lines. **e)** TRF analysis of early passage HCT116 wild type, *MUS81^-/-^*, *MCM10^+/-^* and double mutant cell lines. Yellow dots indicate the location of peak intensity. **f)** *<ι>β</i>*-gal activity expressed as arbitrary fluorescence units normalized to total protein for *MUS81^-/-^* (black) and *MUS81^-/-^*, *MCM10^+/-^* mutant cell lines (gray). Average levels for HCT116 wild type and *MCM10^+/-^* cell lines from Supplementary Figure 2d are indicated with dashed lines, n = 3 replicate wells for all data points. Error bars in **b, d** and **f** indicate standard deviation and significance was calculated using two-tailed students *t-test* with *<.05; **<.01, ***<.001. **g)** Model of *MCM10*-associated telomeropathies. Different cell lineages have an inherent developmental threshold Mcm10-level to achieve complete development and differentiation. As mono- or bi-allelic mutations decrease the amount of functional Mcm10, telomere erosion is accelerated. When Mcm10 function is reduced below the required threshold, eroded telomeres cause replicative senescence and prevent complete development.

Next, we utilized TRF analysis to examine alterations in telomere maintenance. *MUS81* knockout caused telomere erosion, and loss of Mus81 in *MCM10^+/-^* cells caused more significant erosion than in either single mutant. Furthermore, we found elevated *<ι>β</i>*-gal signal in *MUS81* knockouts, with the highest levels detected in the double mutants (Fig. 7f). Taken together, our data clearly demonstrate a requirement for Mus81 in promoting cell proliferation and viability in Mcm10-deficient cells, likely through stimulating replication restart in hard-to-replicate regions, including telomeres.

## DISCUSSION

### Mcm10 deficiency causes genome instability and impairs telomere maintenance

We have demonstrated that *MCM10* is haploinsufficient in HCT116 and RPE-1 cell lines. Whereas both cell types showed a significant reduction in proliferation rate (Fig. 1c, 4b), a global DNA synthesis defect was observed only in HCT116 mutants (Fig. 2f, 4h). The absence of a synthesis defect in RPE-1 mutants is attributable, at least in part, to cell-type specific changes in the replication program during oncogenic transformation of HCT116 cells, as evidenced by the significantly higher levels of origin licensing and DNA synthesis (Fig. 4i-j). It is noteworthy that Mcm10 deficiency caused increased cell death regardless of cell type (Fig. 1i 4c) and that this chronic replication stress was severe enough to stimulate spontaneous reversion of the mutation and rescue of *MCM10* mutant phenotypes in HCT116 cells (Fig. 3f-j, Supplemental Fig. 4). Our analyses of HCT116 mutants also demonstrated that *CDC45* and *MCM4* are haploinsufficient, but suggested that a role in telomere maintenance is unique to *MCM10* (Supplemental Fig. 3).

There is growing consensus that Mcm10 promotes replisome stability^7, 9, 10^, although the mechanism has remained unclear. A recent study demonstrated that yeast Mcm10 is important for bypassing lagging strand blocks^8^, suggesting that Mcm10 is critical when the replisome encounters barriers to CMG translocation. Recently, yeast Mcm10 was shown to prevent translocase-mediated fork regression^34^. If human Mcm10 functions similarly, it might not only prevent stalling but also inhibit excessive fork regression in order to promote restart. In support of this model, we argue that t-complex DNA in *MCM10* mutants was produced by regression of stalled telomeric replication forks. Without sufficient Mcm10, cells must rely on alternative pathways to restart DNA replication. Our data implicate SSEs, including the Mus81 protein, in processing these forks to rescue DNA synthesis and promote *MCM10^+/-^* cell viability (Fig. 7).

Telomere erosion in *MCM10* mutants was not due to decreased telomerase enzymatic function (Fig. 3c, 5d), but to the limiting effect of Mcm10 deficiency. This effect was overcome with sufficiently high telomerase expression (Fig. 5c), although the underlying replication defects persisted. We propose that fork stalling prevented the replisome from reaching chromosome ends; thus, causing telomere shortening through several potential mechanisms. First, telomere replication defects might delay the generation of chromosome ends suitable for telomerase extension. Because telomere replication and telomerase-dependent synthesis primarily occur later in S-phase^35–37^, the replication defect could delay telomere maturation and shorten the cell cycle window when telomerase can act. Second, because reversed replication forks are an aberrant substrate for telomerase^38^, increased fork stalling and regression could impair normal telomerase recruitment or activity. Third, terminal fork arrest in the proximal telomere requiring nucleolytic cleavage to form a functional chromosome end could result in the loss of a significant portion of the distal telomere.

### Development of distinct cell lineages is impaired by *MCM10*-associated telomeropathy

Prior investigations of *MCM10*-associated disease have primarily focused on cancer development. Many studies have described *MCM10* overexpression in a variety of cancer types, wherein the extent of upregulation corresponded with tumor progression and negative clinical outcomes^10^. Presumably, increased Mcm10 expression served to prevent DNA damage and maintain genome stability in cells driven to proliferate. In addition, our data argue that telomere length maintenance in transformed cells, a widely accepted hallmark of cancer, relies on Mcm10^39^. This increased demand in cancer cells also suggests that compounds inhibiting Mcm10 might be useful to preferentially sensitize cancer cells to treatment with common chemotherapeutic drugs or telomerase inhibitors^40^.

We have previously reported one *MCM10-*associated case of NKD^12^. These mutations produced phenotypes similar to the genomic instability caused by NKD-associated mutations in replication factors *MCM4* and *GINS1*^41–43^. It is currently unknown whether these *MCM4* or *GINS1* patient mutations affect telomere maintenance. However, another NKD patient was identified carrying a mutation in *RTEL1* (*regulator of telomere length 1*)^43, 44^, an essential DNA helicase required for the replication of mammalian telomeres^23^. Interestingly, Mcm10 was identified as an RTEL1-interacting protein in mass spectrometry analyses of murine cells^45^, supporting the hypothesis that telomeropathy might underlie NKD. Limited evidence suggests that NK cells have shorter telomeres than T- and B-cells, although they arise from a common progenitor, and that telomerase activity decreases as NK cells differentiate^46–48^. These features may explain why the NK cell lineage is particularly sensitive to defects in telomere replication.

Here, we report *MCM10-*associated RCM with lymphoreticular hypoplasia caused by bi-allelic *MCM10* mutations that resulted in a remarkably similar disease phenotype in three affected patients. To our knowledge, this represents the first connection between alterations in core replisome function and inherited cardiomyopathy. However, cardiomyopathy has been linked with defective telomere maintenance. For example, recent studies have argued that telomere shortening is a hallmark of inherited cardiomyopathies and that long telomeres may be cardioprotective^49, 50^. Additionally, cardiomyopathy is a well-recognized feature of the telomeropathy dyskeratosis congenita (DKC)^51–53^. Intriguingly, mutations in *RTEL1* – discussed above in the context of NKD – have also been strongly linked with telomeropathies including DKC^54, 55^, demonstrating that mutations in the same gene can give rise to clinically distinct pathologies. We propose that this is also the case for *MCM10* and that telomere erosion due to Mcm10 deficiency was the cause of cardiomyopathy in these patients.

The absence of *MCM10-*associated pathology in heterozygous patient family members implies that the disease phenotypes are not attributable to haploinsufficiency. Because each heterozygous combination included one null allele, the hypomorphic alleles in each pair seem necessary to further reduce Mcm10 function and elicit disease. Whereas the NKD-associated missense allele caused a single amino acid substitution, the RCM-associated splice donor mutation is presumably more deleterious, causing a 256 amino acid deletion affecting the major Mcm10 DNA-binding domain^10^ (Fig. 1a, Supplemental fig. 1b,c). It is striking that both compound heterozygous *MCM10* mutations caused immune system abnormalities, albeit to different extents. Furthermore, the more severe combination of alleles (c.236delG;764+5G>A) also impaired cardiac development and is believed to have been lethal *in utero*. In contrast, the NKD-associated alleles (c.1276C>T;1744C>T) resulted in a live-born infant with no overt cardiac phenotype. Based on these observations, we propose that the threshold for Mcm10 function necessary for normal tissue development is cell lineage specific and that differences in the severity of *MCM10* mutations explain why patients presented with distinct but overlapping pathologies (Fig. 7g). Within this model of *MCM10*-associated telomeropathy one key factor that determines the Mcm10 threshold, which still allows for normal differentiation is the rate of telomere shortening inherent to the normal development of a specific cell lineage. Therefore, different cell lineages have their own intrinsic thresholds, explaining the cell type-specific clinical presentations.

## METHODS

### Patient participation

Details of the consent process and qualitative analysis of decision-making in the GM-MDT have been previously described including how, dependent on consent, patients had the option to received “secondary findings”^56^. Patients participated under the Molecular Genetic Analysis and Clinical studies of Individuals and Families at Risk of Genetic Disease (MGAC) protocol approved by West Midlands Research Ethics Committee, reference number 13/WM/0466.

### Clinical samples

Following written informed consent for genetic testing from the patient and/or their parent, blood or post mortem spenic tissue was obtained for genomic DNA extraction. Clinical samples were processed and sequencing results were validated in the Oxford Regional Clinical Molecular Genetic Laboratory. Exome sequencing was performed at the Wellcome Centre for Human Genetics, Oxford. Whole genome sequencing was performed in the BRC/NHS Molecular Diagnostics Laboratory of the Oxford University Hospitals Trust.

### Exome and whole genome sequencing, and bioinformatic analyses

Exome sequencing was performed as described^57^. DNA libraries were prepared from 3 µg patient DNA extracted from whole blood (parents) or spleen (proband). Exome capture was performed using SeqCap EZ Human Exome Library v2.0 (NimbleGen), according to the manufacturer’s instructions, and sequenced using a 100 bp paired-end read protocol on the HiSeq2500 (Illumina). Exome sequence reads were mapped to the hs37d5 reference genome with Stampy^58^. Variants were called with Platypus version 0.5.2^59^. Variants were annotated and analyzed using VariantStudio version 2.2 (Illumina) and Ingenuity VA (Qiagen). Initial data analysis focused on variants associated with an *in silico* panel of 53 cardiomyopathy associated genes, but was later extend to all genes.

Whole genome sequencing was performed in the HICF2 project as described^60, 61^. DNA libraries were prepared from 3 µg patient DNA extracted from spleen. Sequencing was performed using a 100 bp paired-end read protocol on the HiSeq2500 (Illumina). Sequence reads were mapped to the hs37d5 reference genome with Stampy^58^. Variants were called with Platypus version 0.5.2^59^. Variants were annotated and analyzed using VariantStudio version 2.2 (Illumina) and Ingenuity VA (Qiagen). Variant analysis focused on autosomal recessive and X-linked modes of inheritance. For copy-number detection Log_2_R values were generated from bam files and were analyzed together with B-allele frequency outputs. Events were flagged and visualized using Nexus Discovery Edition (BioDiscovery, Inc., El Segundo, CA). The *MCM10* variants were independently validated by Sanger sequencing using BigDye Terminator kit 3.1 (Applied Biosystems) combined with purification using the Agencourt CleanSEQ system. Capillary electrophoresis was performed using an ABI Prism 3730 Genetic Analyser (Applied Biosystems). Effects of the c.764+5G>A variant were assessed using RT-PCR with standard procedures and PCR primers positioned within *MCM10* exons 3 and 8.

### Protein expression and purification

The coding sequence for human Mcm10 internal domain (ID) spanning Ser236 to Gly435, with a preceding HRV 3C protease recognition sequence (LEVLFQGP), was inserted into the NdeI/BamHI sites of the pET28a bacterial expression vector. To mimic the internal deletion of exon 6, a construct lacking the N-terminal 20 amino acid residues (Ser236 to Arg255) was generated and referred to as ΔN20. Recombinant wild-type and mutant (ΔN20) hMcm10-ID were expressed in *E. coli* strain BL21(DE3) and purified essentially as described^40^. The total bacterial lysate was centrifuged at 48,384 x g at 4 °C for 1 hr (Beckman JA-25.50 rotor), and the supernatant was applied to a 5 mL HisPur Ni-NTA resin (Thermo Fisher Scientific) equilibrated with 20 mM Tris-HCl, pH 7.4, 500 mM NaCl, 5 mM β-mercaptoethanol, 5 mM imidazole. Bound His-tagged proteins were subsequently eluted with a linear imidazole concentration gradient up to 300 mM. The eluted proteins were then treated overnight with recombinant HRV 3C protease to remove the N-terminal His-tag and further purified via size exclusion chromatography on a HiLoad 26/600 Superdex 75 column (GE Healthcare) operating with 20 mM Tris-HCl, pH 7.4, 500 mM NaCl, 5 mM β-mercaptoethanol. Monomeric protein peaks were pooled and concentrated using an ultrafiltration device (Amicon) to 24.6 and 15.6 mg ml^-1^ for wild type and ΔN20, respectively, flash cooled by liquid nitrogen in small aliquots and stored at −80°C. Protein concentrations were determined based on UV absorbance measured on a Nanodrop 8000 spectrophotometer and theoretical extinction coefficient values calculated from the protein sequences. The purified proteins after HRV 3C protease cleavage have two extra amino acids (Gly-Pro) on the N-terminus.

### Analytical size-exclusion chromatography

Wild type or ΔN20 hMcm10-ID (200 µL at 5 mg ml^-1^) was injected into Superdex 75 Increase 10/300 GL column (GE Healthcare) operating with the elution buffer containing 20 mM Tris-HCl, pH 7.4, 500 mM NaCl, 5 mM β-mercaptoethanol and a flow rate of 0.4 mL min^-1^. The proteins were detected by UV absorption at 280 nm. To calibrate the column we injected the molecular weight standard (Bio-Rad) including thyroglobulin, *<ι>γ</i>*-globulin, ovalbumin, myoglobin, and vitamin B12 under the same elution condition. All experiments were performed at 4 °C.

### Circular dichroism spectroscopy

Circular dichroism (CD) spectra were collected on a Jasco J-815 CD spectropolarimeter with a temperature controller, using quartz cuvettes with the pathlength of 0.1 cm (Starna Ltd.). The samples contained purified protein at 0.1 mg mL^-1^ in 50 mM sodium phosphate, pH 7.4 and 0.5 mM TCEP. The blank (baseline) sample contained the same buffer without protein. Data acquisition was performed for the wavelength range of 190-260 nm with 1 nm steps, at the temperatures of 25, 53, 60, 65, 70, 75, and 80 °C (only the data for 25 and 60 °C are shown). Fresh protein sample was prepared for the measurement at each temperature and the spectra were scanned five times. The data are presented as molar ellipticity ([q] / deg cm² dmol^-1^ x 10^5^) plotted versus wavelength. To monitor thermal denaturation, CD spectra for the wavelength range from 205 to 220 nm, in 1 nm steps, were collected over the temperature range from 40 to 90 °C in 1 °C increments. The spectra were scanned five times at each temperature following a 10-sec equilibration period. Single wavelength melt curves were obtained by plotting the measured CD signal at 205 nm against temperature. The curves were normalized by setting the minimum (greatest magnitude) ellipticity at 40 °C as 0 and maximum ellipticity at 90 °C as 1. The temperature at which the normalized CD signal reaches 0.5 was defined as the melting temperature (T_m_).

### Cell lines

HCT116 cells were grown in McCoy’s 5A medium (Corning 10-050-CV) supplemented with 10% FBS (Sigma F4135), 1% Pen Strep (Gibco 15140) and 1% L-Glutamine (Gibco 205030). hTERT RPE-1 cells were grown in DMEM/F12 medium (Gibco 11320) supplemented with 10% FBS and 1% Pen Strep. Cells were cultured at 37°C and 5% CO_2_.

### Cell line generation using rAAV

HCT116 *MCM10^+/-^* (exon 14) cell lines were generated using rAAV-mediated gene targeting^62^. The conditional vector pAAV-MCM10-cond was constructed using Golden Gate cloning and designed as described^62^. The MCM10-cond rAAV was generated by co-transfection of pAAV-MCM10-cond, pAAV-RC^62^, and pHelper^62^ into HEK293 cells using Lipofectamine LTX (Invitrogen 15338030) following standard protocols^62^. The first round of targeting replaced *MCM10* exon 14 with a wild type allele and a downstream neomycin (G418) selection cassette both flanked by *loxP* sites (“floxed”). Targeted clones were selected using 0.5 mg/ml G418 (Geneticin G5005). Resistant clones were screened by PCR using primers within the neomycin cassette and outside the rAAV homology arms to confirm locus-specific targeting, and Cre recombinase transiently expressed from an adenoviral vector (AdCre; Vector Biolabs #1045) was then used to remove the neomycin selection cassette as described^62^. The second round of *MCM10* gene targeting used the same rAAV vector and replaced the wild type allele with a floxed allele and a downstream floxed neomycin selection cassette. G418-resistant clones were screened by PCR to confirm locus-specific targeting. AdCre recombinase was then used to remove the neomycin selection cassette and resulted in the generation of heterozygous *MCM10* clones. The *MCM10* exon 14 genotype was subsequently screened and confirmed using primers flanking the exon.

### Cell line generation using CRISPR/Cas9

HCT116 and RPE-1 *MCM10^+/-^* exon 3, HCT116 *CDC45^+/-^* exon 3 and HCT116 *MCM4^+/-^* exon 2 cell lines were generated using CRISPR/Cas9 gene targeting. Guide RNAs (gRNAs) were cloned into a CRISPR/Cas9 plasmid hSpCas9(BB)-2A-GFP (PX458; Addgene #48138)^63^. Cells were transfected with CRISPR/Cas9 plasmid containing gRNA using the Neon Transfection System (Invitrogen MPK5000) following standard protocols. Two days post-transfection GFP-positive cells were collected by flow cytometry. Subcloned cells were screened for correct targeting by PCR amplification and restriction enzyme digestion (*MCM10* exon 3, Hpy199III (NEB R0622); *CDC45* exon 3, PflMI (NEB R0509), XcmI (NEB R0533) or AlwNI (NEB R0514); *MCM4* exon 2, BglI (NEB R0143). Specific mutations were identified by PCR and Illumina sequencing or PCR, DNA sequencing and TIDE (<u>T</u>racking of Indels by <u>De</u>composition) analyses^64^.

To generate HCT116 *MUS81^-/-^* exon 2 mutant cell lines, each parental line was transfected with 1 µL of 100 µM sgRNA (Synthego Corporation) and 1 µg Cas9 mRNA (TriLink #L-7206) using the Neon Transfection System following standard protocols. Three days post-transfection cells were subcloned. Subclones were screened for correct targeting by PCR amplification, DNA sequencing and TIDE analyses.

### Cell line generation using plasmid transfection

To generate HCT116 cells lines expressing super-telomerase^31^, pBABEpuroUThTERT+U3-hTR-500^65^ (Addgene #27665) was linearized with restriction enzyme ScaI (NEB R3122) and transfected into wild type or mutant HCT116 cell lines following standard Lipofectamine 3000 protocols (Invitrogen L3000). Stable cell lines were generated using puromycin selection (1 µg/ml; Sigma P7255) followed by subcloning.

### Cell proliferation

Cells were plated at 50,000 cells per well (RPE-1) or 100,000-125,000 cells per well (HCT116) in 6-well plates. Cell counts were performed 3-days after seeding using Trypan Blue (Invitrogen T10282) on Countess slides (Invitrogen C10283) using a Countess automated cell counter (Invitrogen C20181).

### Protein extraction, chromatin fractionation and western blotting

For preparation of whole cell extracts, cells were lysed in RIPA (50 mM Tris-HCl, pH 8.0, 150 mM NaCl, 10 mM NaF, 1% NP-40, 0.1% SDS, 0.4 mM EDTA, 0.5% sodium deoxycholate, 10% glycerol) buffer for 10 min and then centrifuged at 16,000 *g* for 10 min. Cleared lysates were collected, mixed with SDS loading buffer and boiled before fractionation by SDS-PAGE and analyses by western blot. Chromatin fractions were isolated as described^66^. Extracts were prepared by lysis in Buffer A (10 mM HEPES pH 7.9, 10 mM KCl, 1.5 mM MgCl_2_, 0.34 M sucrose, 10% glycerol, 0.1% Triton X-100 and protease inhibitors). Insoluble nuclear proteins were isolated by centrifugation and chromatin-bound proteins were subsequently released by sonication. Remaining insoluble factors were cleared by centrifugation before fractionation by SDS-PAGE and western blot analyses. Primary antibodies were incubated in 5% BLOT-QuickBlocker (G-Biosciences 786-011) as follows: rabbit anti-Mcm10 (Bethyl A300-131A; 1:500), rabbit anti-Mcm10 (Novus, H00055388-D01P, 1:500), mouse anti-Cdc45 (Santa Cruz G12, SC55568; 1:500), mouse anti-Mcm4 (Santa Cruz G7, SC28317; 1:500), mouse anti-Mus81 (Abcam ab14387; 1:500), mouse anti-PCNA (Abcam Ab29; 1:3,000), rabbit anti-RPA32 (S4/8) (Bethyl A300-245A; 1:2000), mouse anti-GAPDH (GeneTex GTX627408; 1:5,000), mouse anti-*<ι>α</i>*-Tubulin (Millipore T9026, clone DM1A; 1:10,000). Secondary antibodies were incubated in 5% BLOT-QuickBlocker (G-Biosciences 786-011) at 1:10,000 dilutions, including goat anti-mouse HRP conjugate (Jackson Laboratories 115-035-003), goat anti-rabbit HRP conjugate (Jackson Laboratories 111-035-144), goat anti-mouse HRP conjugate (BioRad, 1706516), donkey anti-rabbit HRP conjugate (Amersham NA9340). Detection was performed using WesternBright Quantum detection kit (K-12042-D20). Quantification was performed using FIJI and Microsoft Excel. Image preparation was performed using Adobe Photoshop.

### FACS analysis

For flow cytometry analyses of cell cycle, DNA synthesis and origin licensing wild type and *MCM10+/-* HCT116 and hTERT RPE-1 cells lines were treated as described^16^. Briefly, cells were incubated with 10 µM EdU (Santa Cruz sc-284628) for 30 min before harvesting with trypsin. Soluble proteins were extracted in CSK (10 mM PIPES pH 7.0, 300 mM sucrose, 100 mM NaCl, 3 mM MgCl_2_ hexahydrate) with 0.5% Triton X-100, then cells were fixed in PBS with 4% PFA (Electron Microscopy Services) for 15 min. Cells were labeled with 1 µM AF647-azide (Life Technologies A10277) in 100 mM ascorbic Acid, 1 mM CuSO_4_, and PBS to detect EdU for 30 min, at room temperature. Cells were washed, then incubated with MCM2 antibody 1:200 (BD Biosciences #610700) in 1% BSA in PBS with 0.5% NP-40 for 1 hr at 37°C. Next, cells were washed and labeled with donkey anti-mouse AF488 secondary antibody 1:1,000 (Jackson Immunoresearch 715-545-150) for 1 hr at 37°C. Lastly, cells were washed and incubated in DAPI (Life Technologies D1306) and 100 ng/mL RNAse A (Sigma R6513) overnight at 4°C. Samples were run on an Attune NxT (Beckman Coulter) or LSR II (BD Biosciences) flow cytometer and analyzed with FCS Express 6 (De Novo Software) or FlowJo v10.6.1 and Microsoft Excel.

For flow cytometry analysis of apoptosis, cells were seeded in 6-well plates (HCT116 wild-type at 150,000 cells/well, HCT116 mutants at 200,000-400,000 cells/well, RPE-1 wild-type at 50,000 cells/well, and RPE-1 mutants at 75,000 cells/well) and allowed to proliferate for approximately 72 hr. Adherent and floating cells were collected, washed with 1x PBS twice, and stained using the APC Annexin V apoptosis detection kit (Biolegend 640932) according to the manufacturer’s instructions. Samples were analyzed on a FACSCanto A V0730042 (BD Biosciences). Apoptotic cells were identified by annexin V staining while cell viability was determined by PI staining. Data was analyzed using FlowJo v10.6.1 and Microsoft Excel.

### DNA combing

HCT116 cells were plated at 1×10^6^ cells per 10 cm plate 48 hr prior to labeling. Cells were incubated with 25 µM IdU (Sigma C6891) for 20 or 30 min, rinsed with pre-warmed medium and then incubated with 200 µM CldU (Sigma I7125) for 30 min. Approximately 250,000 cells were embedded in 0.5% agarose plugs (NuSieve GTG Agarose, Lonza, 50080) and digested for 48 to 72 hr in plug digestion solution (10 mM Tris-HCl, pH 7.5, 1% Sarkosyl, 50 mM EDTA and 2 mg/ml Proteinase K). Plugs were subsequently melted in 50 mM MES pH 5.7 (Calbiochem #475893) and digested overnight with *<ι>β</i>*-agarase (NEB M0392). DNA combing was performed using commercially available coverslips (Genomic Vision COV-001). Integrity of combed DNA for all samples was checked via staining with YOYO-1 (Invitrogen Y3601). Combed coverslips were baked at 60°C for 2 to 4 hr, cooled to room temperature (RT) and stored at −20°C. DNA was denatured in 0.5 M NaOH and 1 M NaCl for 8 min at RT. All antibody staining was performed in 2% BSA in PBS-Triton (0.1%). Primary antibodies include rabbit anti-ssDNA (IBL 18731), mouse anti-BrdU/IdU (BD Biosciences 347580) and rat anti-BrdU/CldU (Abcam ab6326). Secondary antibodies include goat anti-mouse Cy3.5 (Abcam ab6946), goat anti-rat Cy5 (Abcam ab6565) and goat anti-rabbit BV480 (BD Horizon #564879). Imaging was performed using Genomic Vision EasyScan service. Image analyses were blinded and used the Genomic Vision FiberStudio software and data/statistical analyses were performed in Microsoft Excel and GraphPad Prism 8.

For telomere specific analyses DNA replication in HCT116 super-telomerase cell lines, cells were plated as described above. Cells were incubated with 200 µM CldU (Sigma I7125) for 1 hr, rinsed with pre-warmed medium and then grown without label for 2 hr (repeated 3 additional times). Approximately 500,000 cells were embedded per plug and digested as described above. Plugs were next digested with either RsaI (NEB R0167) or HinfI (NEB R0155) restriction enzyme in 1x NEB CutSmart Buffer to degrade non-telomeric DNA. Samples were then melted, combed, baked and stored as described above. DNA was denatured in 0.5 M NaOH and 1 M NaCl for 12 min at RT. Telomeres were detected using TelG-Alexa488-conjugated PNA probe (PNA Bio F1008) with a 1:25 dilution in hybridization solution (35% formamide, 10 mM Tris-HCl pH 7.5, 0.5% blocking buffer pH 7.5 (100 mM maleic acid, 150 mM NaCl, 10% Roche blocking reagent #11096176001). Antibody staining was performed in 3% BSA in PBS-Triton (0.1%). CldU-labeled DNA was detected using primary rat anti-CldU (Abcam ab6326), followed by secondary goat anti-rat AF555 (Invitrogen A21434) antibodies. Imaging was performed using an EVOS FL imaging system (ThermoFisher AMF43000). Image analyses were blinded and used FIJI and Adobe Photoshop. Statistical analysis was performed using Microsoft Excel.

### Clonogenic survival assay

Wild type cells were plated at 500 cells/well, *MCM10^+/-^* cells were plated at 800 or 1000 cells/well unless otherwise noted. After 24 hr, the medium was removed and replaced with drug containing medium (hydroxyurea, Acros Organics 151680250; mitomycin C, Sigma M4287) or cells were exposed to UV (UVP CL1000 crosslinker) and fresh medium was added. Cells were incubated for 12 to 14 days, washed in PBS, fixed in 10% acetic acid/10% methanol and stained with crystal violet. Colonies reaching a minimum size of 50 cells were counted manually and normalized to the average colony number in untreated wells. Statistical analysis was performed using Microsoft Excel. Plates were imaged using an Epson Expression 1680 scanner.

### Cellular senescence assay

HCT116 cells were plated at 2,000 to 4,000 cells per well and RPE-1 cells were plated at 1,000 to 1,5000 cells per well in 96-well plates and allowed to recover for three days. The *<ι>β</i>*-galactosidase activity was measured with the 96-Well Cellular Senescence Assay Kit (Cell Biolabs CBA-231) following the manufacturer’s instructions with the following modifications. Cell lysates were centrifuged in v-bottom 96-well plates rather than microcentrifuge tubes and total protein concentration was determined using Protein Assay Dye (BioRad #500-0006) following standard protocols. Plates were imaged using a VICTOR^3^V 1420 Multilabel Counter (Perkin Elmer). The *<ι>β</i>*-galactosidase activity was normalized to total protein concentration and shown as arbitrary fluorescence units. Analysis and statistical tests were performed using Microsoft Excel.

### Telomere restriction fragment (TRF) Analysis

Genomic DNA was extracted from ∼1 x 10^7^ cells using a modified version of the Gentra Puregene Cell Kit cell extraction protocol (Qiagen 158745). Integrity of genomic DNA and absence of contaminating RNA was confirmed via 1% agarose 1x TAE gel electrophoresis. Subsequently, 30 to 40 μg of genomic DNA was digested with HinfI (NEB R0155) and RsaI (NEB R0167), as described. For each sample, 8 to 12 μg of digested genomic DNA was resolved overnight on a 0.7% agarose 1x TBE gel. Gels were depurinated, denatured, and neutralized, followed by overnight capillary transfer to a Hybond-XL membrane (GE Healthcare RPN303S). Telomere probe was labeled using T4 polynucleotide kinase (NEB M0201) and *<ι>γ</i>*-P^32^-ATP (Perkin Elmer NEG035C) and purified using quick spin columns (Roche 11-273-949-001). Membranes were pre-hybridized for 1 hr with Church buffer at 55°C, then hybridized with a *<ι>γ</i>*-P^32^-end-labeled telomere probe ((C_3_TA_2_)_4_) in Church buffer at 55°C overnight. Membranes were washed three times with 4x SSC and once with 4x SSC + 0.1% SDS, each for 30 min, exposed to a phosphorimaging screen, detected with a Typhoon FLA 9500 imager and processed using FIJI and Adobe Photoshop. For TRF analyses using telomerase inhibitor, a 10 mM stock solution of BIBR1532 (Tocris #2981) in DMSO was prepared and diluted in appropriate growth medium to a final concentration of 10 µM as described^30^.

### Telomere 2D gel analysis

Samples were collected and prepared as described above for TRF analyses. For S1 nuclease digested samples, 20 µg of RsaI/HinfI digested DNA was digested with 100 U of S1 nuclease (Thermo Scientific #EN0321) for 45 min at room temperature. For each sample, 8 to 12 μg of digested genomic DNA was resolved overnight on a 0.5% agarose 1x TBE gel. Sample lanes were cut out, re-cast and resolved on a 1.2% agarose 1x TBE gel. Gels were treated, transferred to Hybond-XL membrane, labeled and imaged as described above for TRF analyses.

### Telomeric repeat amplification protocol (TRAP) assay

TRAP assays were performed following the manufacturer’s protocol for the TRAPeze Telomerase Detection Kit (EMD-Millipore S7700). Briefly, whole cell extracts were prepared using 1x CHAPS lysis buffer and stored at −80 DC. Protein concentration was measured using Protein Assay Dye (BioRad #500-0006) in comparison to a BSA (NEB B9000S) standard curve following standard protocols. TRAP reactions utlized Platinum Taq DNA Polymerase (Invitrogen 10966), and were resolved on 10% polyacrylamide 0.5x TBE gels, stained for 1 hr with SYBR GOLD (Invitrogen S11494) and detected with a Typhoon FLA 9500 imager and processed using FIJI and Adobe Photoshop.

### Immunofluorescence analyses of telomeres

For t-FISH asynchronous HCT116 populations were incubated for 1 hr in 0.25 µg/mL KaryoMAX colcemid (Gibco 15212-012). Cells were collected and washed in 75 mM prewarmed KCl. Cells were then fixed three times in methanol:acetic acid (3:1) by adding fixative solution dropwise with constant gentle agitation by vortex. Following fixation, cells were dropped onto microscope slides and metaphase spreads were allowed to dry overnight. Next, slides were rehydrated in 1x PBS, followed by fixation in 3.7% formaldehyde. Slides were then washed twice in 1x PBS, rinsed in ddH2O, dehydrated in an ethanol series (70%, 85%, 95%) pre-chilled to −20 DC and air dried.

FISH was performed with TelC-Cy3 (PNA Bio F1002) and CENPB-Alexa488 (F3004) warmed to 55°C then diluted 1:300 in hybridization buffer (70% formamide, 10 mM Tris pH 7.4, 4 mM Na_2_HPO_4_, 0.5 mM citric acid, 0.25% TSA blocking reagent (Perkin Elmer FP1012), and 1.25mM MgCl2) preheated to 80 °C. Slides were denatured with probe at 80 °C, then allowed to incubate at room temperature in a humid chamber for 2 hr. Next, slides were washed twice in PNA wash A (70% formamide, 0.1% BSA, 10 mM Tris pH 7.2) and three times in PNA wash B (100 mM Tris pH 7.2, 150 mM NaCl, 0.1% Tween-20). The second PNA wash B contained DAPI (Life Technologies D1306) at a 1:1000 concentration. Slides were then dehydrated and dried as described above prior to mounting. Slides were blinded prior to imaging and captured using a Zeiss Spinning Disk confocal microscope. Image analyses were blinded and used FIJI and Adobe Photoshop. Statistical analysis was performed using Microsoft Excel.

For analyses of telomere sister chromatid exchange, HCT116 ST cells were cultured in the presence of BrdU:BrdC (final concentration of 7.5 mM BrdU (MP Biomedicals 100166) and 2.5 mM BrdC (Sigma B5002)) for 12 hr prior to harvesting. KaryoMAX colcemid (Gibco 15212-012) was added at a concentration of 0.1 μg/mL during the last two hours. CO-FISH was performed as described^67^ using a TelC-Alexa488-conjugated PNA probe (PNA Bio F1004) followed by a TelG-Cy3-conjugated PNA probe (PNA Bio F1006). Images were captured using a Zeiss Spinning Disk confocal microscope. Image analyses were blinded and used FIJI and Adobe Photoshop. Statistical analysis was performed using Microsoft Excel.

### Statistical analysis

Statistical details of experiments including the statistical tests used, value of *n*, what *n* represents, dispersion and precision measures (e.g. average, standard deviation), as well as how significance was defined is indicated in each figure and corresponding figure legend. The software used for statistical analysis of each type of experiment is indicated in the corresponding Method Details section.

### Data availability

The authors declare that all data related to the findings of this study are available within the article and supplementary information, or are available from the corresponding author upon reasonable request.

## Acknowledgments

We thank members of the Bielinsky laboratory for helpful discussions. We also thank L. Harrington for generously sharing reagents and R. Rebbeck for help with CD experiments. This work was supported by NIH grants GM074917 (A.K.B.), GM134681 (A.K.B.), GM083024 (J.G.C.), GM102413 (J.G.C.), CA190492 (E.A.H.), GM118047 (H.A.) and T32-CA009138 (R.M.B., W.L., C.B.R.), by NSF Fellowship DGE-1144081 (J.P.M.), a UNC Dissertation Completion Fellowship (J.P.M.), NIH National Center for Advancing Translational Sciences grants TL1R002493 and UL1TR002494 (M.M.S.) and the ARCS Foundation (C.B.R.). We wish to acknowledge the University of Minnesota Flow Cytometry Resource, the University of Minnesota Imaging Centers, the University of Minnesota Genomics Center and the Masonic Cancer Center Cancer Genomics Shared Resource supported by P30 CA077598. The University of North Carolina Flow Cytometry Core Facility is supported in part by P30 CA016086. Exome sequencing was supported in part by the National Institute for Health Research Oxford Biomedical Research Centre Program. Genome sequencing was conducted as part of an independent research program commissioned by the Health Innovation Challenge Fund (R6-388 / WT 100127), a parallel funding partnership between the Wellcome Trust and the Department of Health. The views expressed in this publication are those of the authors and not necessarily those of the Wellcome Trust or the Department of Health.

## Author contributions

R.M.B. designed and performed experiments, analyzed data, prepared the figures and co-wrote the manuscript. W.L., J.P.M., M.K.O., M.M.S., L.Y., J.T., A.T.P., J.C.T., H.D., J.C., E.O., H.W. and E.B. performed experiments and analyzed data. G.S. helped with data interpretation. L.W., C.B.R., D.B., J.H., and A.H. performed experiments. A.K.B. designed experiments, supervised the study and co-wrote the manuscript. All authors reviewed and edited the manuscript.

## Competing interests

The authors declare no competing interests.

## Materials and correspondence

Correspondence and materials requests should be addressed to Anja-Katrin Bielinsky

## SUPPLEMENTARY INFORMATION

### Supplementary Note

#### Restrictive cardiomyopathy with lymphoreticular hypoplasia

The proband was a male fetus of a gravida 2 para 1 healthy Caucasian female. There was no familial consanguinity. A routine antenatal scan at 12 weeks gestation was normal. Routine antenatal scanning at 20 weeks gestation revealed a pericardial effusion. Follow up scans showed progressive fetal hydrops and reduced cardiac function, with cardiac morphology suggestive of a restrictive cardiomyopathy (RCM). Intra-uterine death was recorded at 27 weeks gestation and the fetus was delivered. Karyotyping on amniotic fluid was normal. A post mortem examination was undertaken. No dysmorphic facial features, limb abnormalities or major structural cardiac abnormalities were noted. However, the right atrium (RA) was severely dilated and the left ventricle (LV) was thickened. Myocardial histology was non-specific but showed enlarged myofiber nuclear size and endocardial fibroelastosis consistent with a cardiomyopathy. The thymus and spleen were hypoplastic, weighing 0.8 g (normal range for gestation 2.04 +/- 0.39 g) and 0.82 g (normal range for gestation 1.24 +/- 0.36 g), respectively. Histology of the spleen showed autolyzed tissue with no specific pathological features. Further molecular genetic investigation including a 57 gene “cardiomyopathy panel”, mitochondrial genetic analysis and oligonucleotide array comparative genomic hybridization (CGH) revealed no plausible pathogenic variant.

The next pregnancy underwent detailed antenatal ultrasound scans at 9 and 12 weeks of gestation that were normal. A scan at 16 weeks revealed ascites with pleural and pericardial effusions. Cardiac function was reduced. Termination of pregnancy was carried out at 17 weeks gestation. No dysmorphic features were noted. There were no limb abnormalities. The RA was severely dilated and the LV was larger than the RV, but no structural congenital cardiac defect was identified. Myocardial histology revealed enlarged myofiber nuclei with endomyocardial fibroelastosis. The thymus and spleen were hypoplastic, weighing 0.04 g (normal range for gestation 0.18 +/- 0.06 g) and 0.05 g (normal range for gestation 0.14 +/- 0.07 g), respectively. Histology of both was unremarkable.

The third affected pregnancy followed a remarkably similar course to those described above with an antenatal diagnosis of RCM with fetal hydrops. Termination of pregnancy was undertaken. The RA was enlarged but no structural heart disease was noted. Myocardial histology revealed myofiber nuclear hypertrophy and endomyocardial fibroelastosis, consistent with a diagnosis of cardiomyopathy. The thymus and spleen were hypoplastic, weighing 0.32 g (normal range for gestation 1.71 +/- 0.54 g) and 0.69 g (normal range for gestation 1.05 +/- 0.25 g), respectively. The parents and three other clinically unaffected siblings had a normal ECG and echocardiogram.

#### Exome and Genome Sequencing of RCM patients and family

Exome sequencing of the parents and proband yielded no strong candidate variants from a panel of known cardiomyopathy associated genes. A gene agnostic trio analysis was undertaken. This generated an extensive list of variants of uncertain significance. To reduce the number of candidates and/or identify additional variants the third affected fetus underwent genome sequencing and this data was analyzed together with the original three exomes. Given the pedigree structure, analysis focused on autosomal recessive and X-linked modes of inheritance. Compound heterozygous variants c.[236delG];[764+5G>A] in *MCM10* (NM_018518.5; Figure 1a) remained plausible candidate variants following this quad analysis. These variants were validated using Sanger sequencing and were shown to segregate within the family in a manner consistent with them being autosomal recessive pathogenic variants (Figure 1b). Each unaffected individual had at least one wild type *MCM10* allele.

### Supplementary figure legends

**Supplementary Fig. 1.**
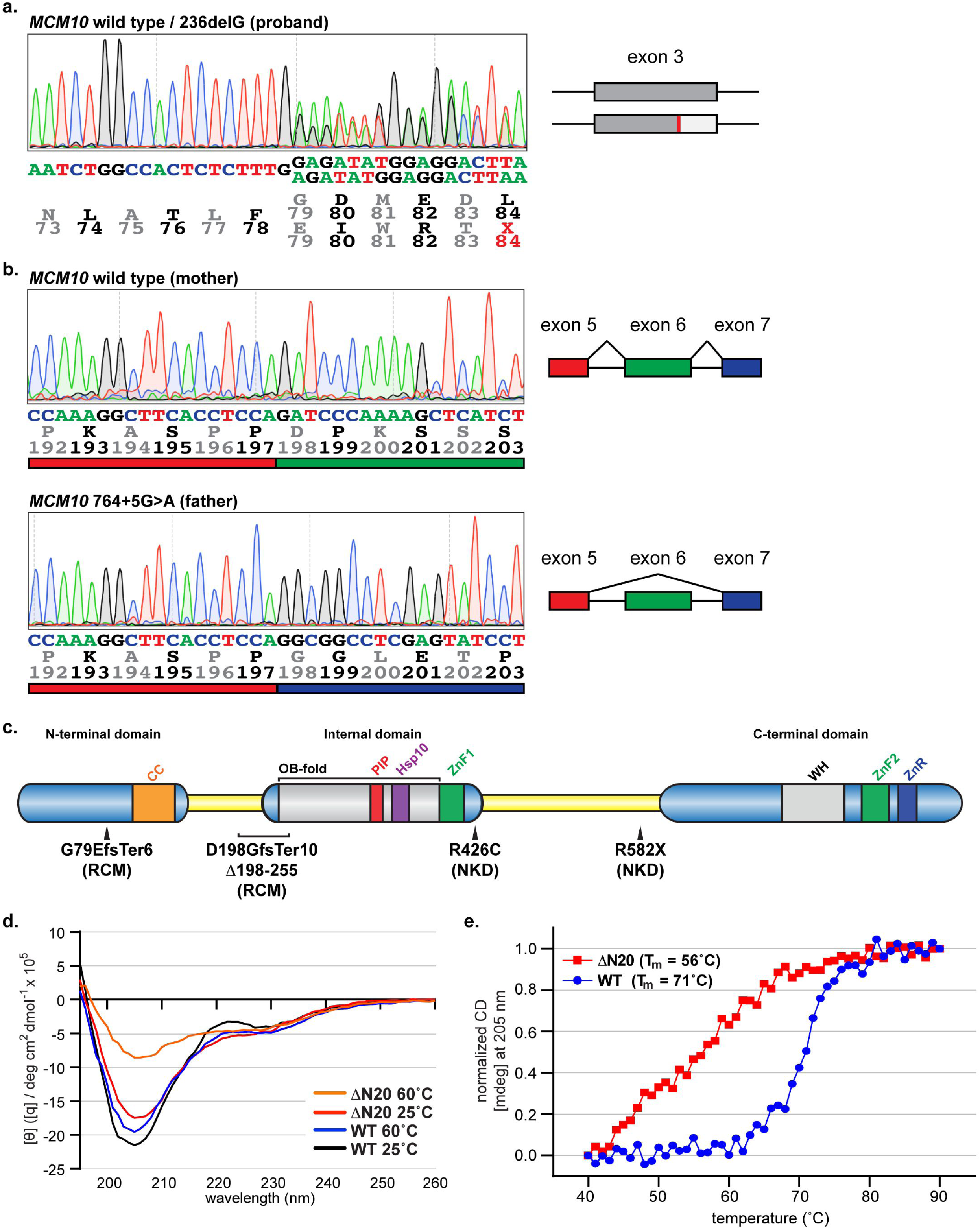
Identification and modeling of *MCM10* patient mutations. **a)** Sequencing analysis from the proband validating the c.236delG mutation in *MCM10* exon 3 that introduces a premature stop codon. Sequencing base calls and corresponding translation is shown below the trace image. On the 3’ end the base calls the wild type allele is shown on top and the mutant allele with a single G deletion is shown on the bottom. The translation of each codon is shown with the single letter abbreviations for each amino acid and their position in the Mcm10 protein. Annotation is in reference to *MCM10* transcript NM_018518.5. (Right) Cartoon depiction of the wild type and mutant c.236delG exon 3 *MCM10* alleles. **b)** Sequencing analysis from the mother showing the wild type *MCM10* exon5/6 junction and from the father validating the c.764+5G>A mutation that results in loss of exon 6 from the mature mRNA. Sequencing base calls and corresponding translation is shown below each trace image. The translation of each codon is shown with the single letter abbreviations for each amino acid and their position in the Mcm10 protein. Below each translation is a bar indicating whether that region is coded for by *MCM10* exon 5 (red), exon 6 (green) or exon 7 (blue). Annotation is in reference to *MCM10* transcript NM_018518.5. (Right) Cartoon depiction of the wild type and mutant c.764+5G>A alleles. **c)** Modified cartoon of human Mcm10^70^ including the N-terminal domain that harbors a coiled-coil (CC, orange) motif. The internal domain contains a PCNA-interacting peptide (PIP) box (red), Hsp10-like domain (purple), an oligonucleotide/oligosaccharide binding (OB)-fold (gray) and zinc-finger motif 1 (ZnF1, green). The C-terminal domain contains ZnF2 (green), the zinc ribbon (ZnR, blue) and winged helix motif (WH, light gray). Mcm10 structural domains are connected by two flexible linker regions (yellow). The location of the RCM-associated G79EfsTer6 and D198GfsTer10 (*<ι>Δ</i>*198-255) and NKD- associated R426C and R582X mutations are indicated. Annotation is in reference to *MCM10* transcript NM_018518.5. **d)** CD spectra comparing WT and *<ι>Δ</i>*N20 Mcm10-ID at 25°C and 60°C. **e)** Melt curves for WT and *<ι>Δ</i>*N20 Mcm10-ID obtained by plotting the normalized CD signal (ellipticity) at 205 nm from 40°C to 90°C.

**Supplementary Fig. 2.**
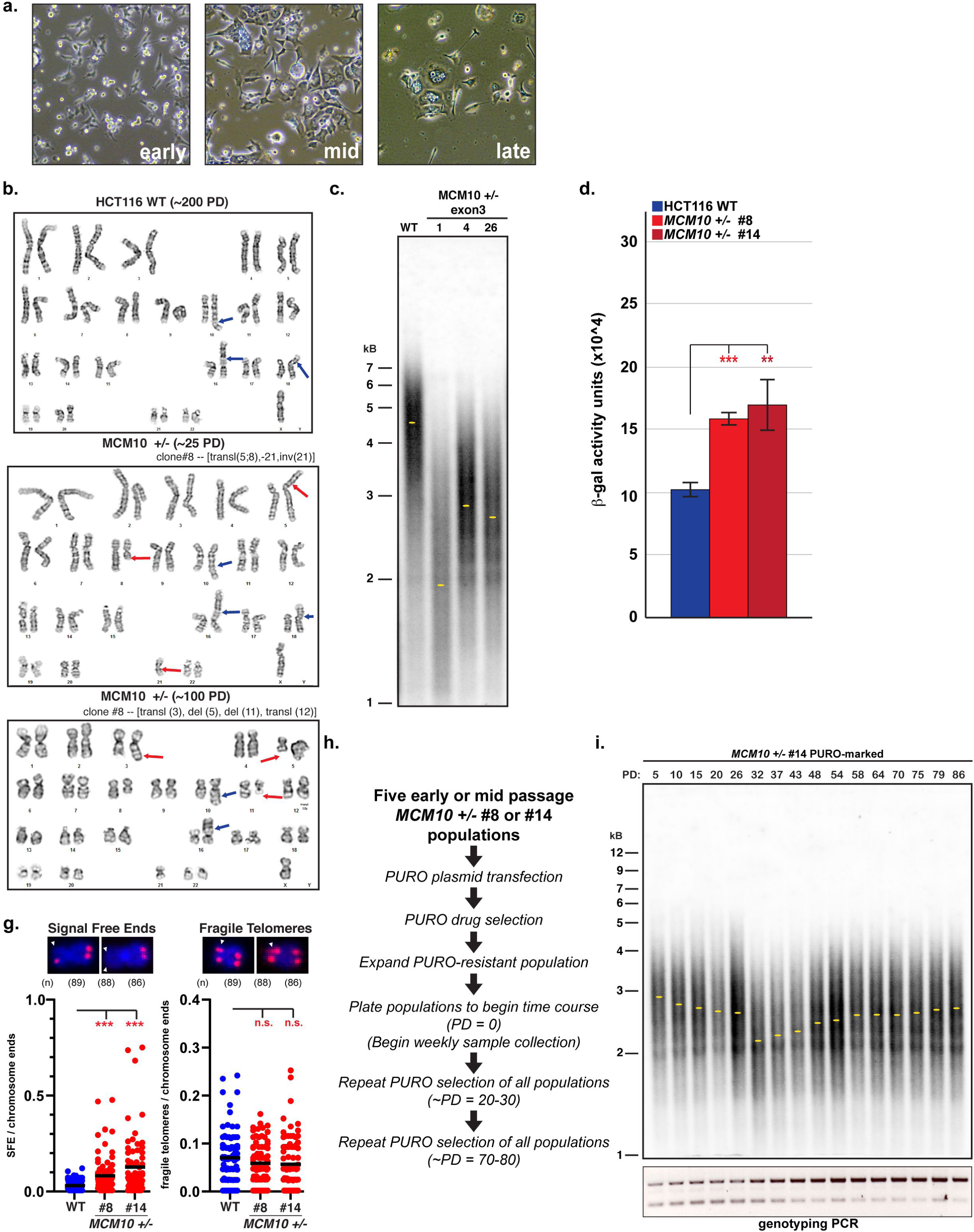
Characterization of genomic instability in *MCM10^+/-^* cell populations. **a)** Representative phase contrast images of early, middle and late passage *MCM10^+/-^* cell populations. **b)** Example karyotypes from late passage HCT116 wild type (top) and mid-passage (middle) or late passage (bottom) *MCM10^+/-^* cells. Blue arrows indicate expected HCT116 genomic aberrations. Red arrows indicate non-clonal genomic aberrations. **c)** TRF analysis comparing early passage HCT116 wild type cells with clonal Mcm10- deficient populations carrying inactivating mutations in one copy of *MCM10* exon 3. Yellow dots indicate the location of peak intensity. **d)** Quantification of *<ι>β</i>*-gal activity expressed as arbitrary fluorescence units normalized to total protein for HCT116 wild type (blue) and clonal *MCM10^+/-^* cell lines (red). Error bars indicate standard deviation and statistical significance was calculated using two tailed student’s *t-test* with *<.05; **<.01, ***<.001; n=3 replicate wells for all data points. **g)** Quantification of signal free-ends (left) and fragile telomeres (right) in late passage HCT116 wild type (blue) and *MCM10^+/-^* cells (red). Error bars indicate standard deviation and statistical significance was calculated using two-tailed student’s *t-test* with ***<.001; n = number of metaphases quantified per cell line. **h)** Flow chart for experiment to generate and analyze PURO-marked *MCM10^+/-^* cell lines. **i)** Analysis of a PURO-marked HCT116 *MCM10^+/-^* het #14 cell population tracking average telomere length by TRF (top) and reversion of the exon 14 locus by PCR (bottom). PDs at each time point is indicated. Yellow dots indicate the location of peak intensity.

**Supplementary Fig. 3.**
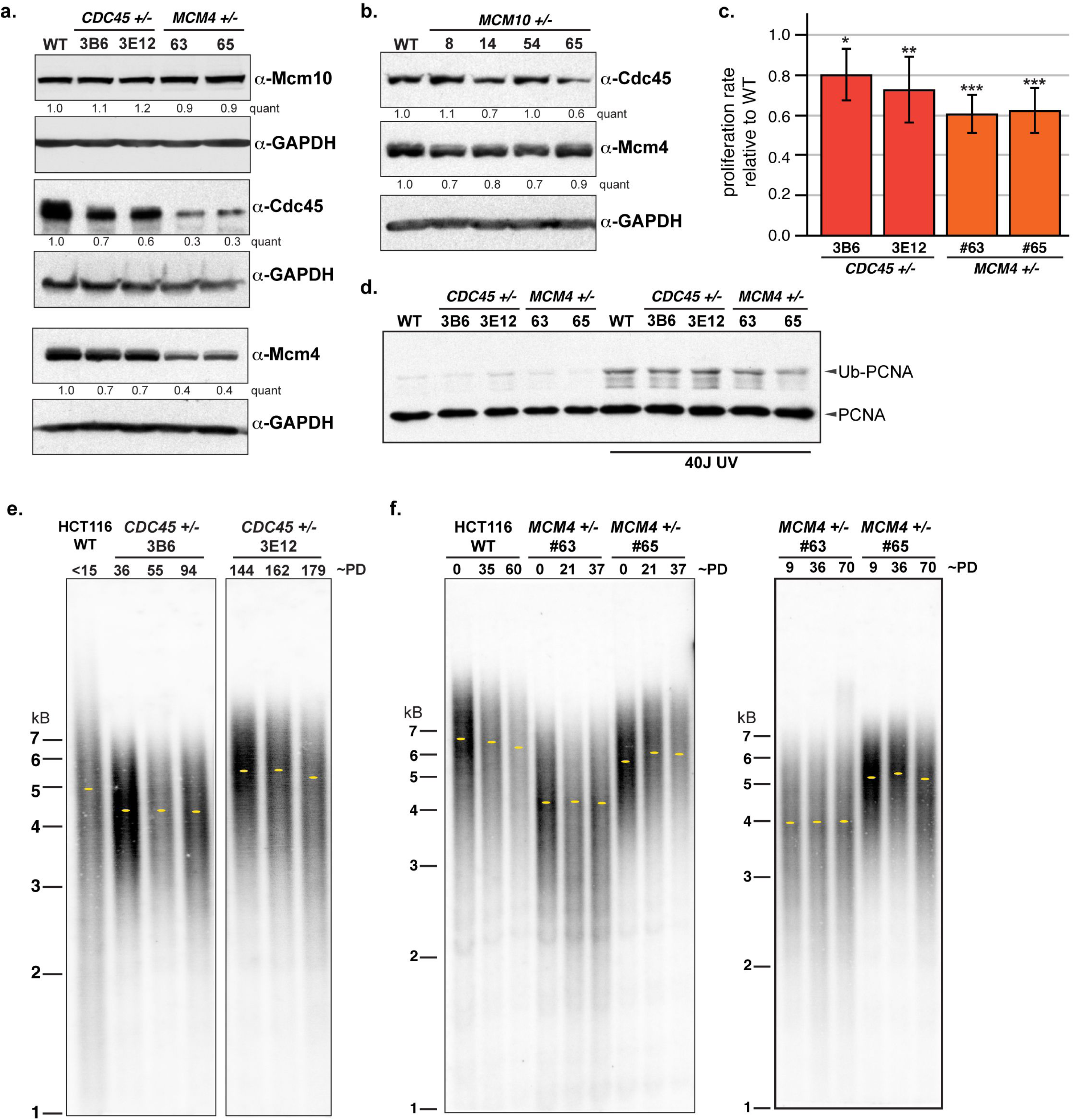
*CDC45* and *MCM4* deficiency causes haploinsufficiency but not telomere erosion in HCT116 cell lines. **a)** Western blot analyses for Mcm10, Cdc45 and Mcm4, with GAPDH loading controls. Quantification of Mcm10, Cdc45 or Mcm4 levels normalized to loading control, relative to the first lane wild-type sample is indicated. **b)** Western blot analyses for Cdc45 or Mcm4 with GAPDH as a loading control. Quantification of Cdc45 or Mcm4 levels normalized to loading control, relative to the first lane wild-type sample is indicated. **c)** Quantification of growth rate in *CDC45^+/-^* (red) and *MCM4^+/-^* (orange) cell lines normalized to HCT116 wild type cells. For each cell line n=6 replicate wells across three biological replicates; error bars indicate standard deviation and statistical significance was calculated using two-tailed student’s *t-test* with *<.05; **<.01, ***<.001. **d)** TRF analysis in HCT116 wild type and *CDC45^+/-^* cell lines. Estimated PDs is indicated. Yellow dots indicate the location of peak intensity. **e)** TRF analysis in HCT116 wild type and *MCM4^+/-^* cell lines. Estimated PDs is indicated. Yellow dots indicate the location of peak intensity.

**Supplementary Fig. 4.**
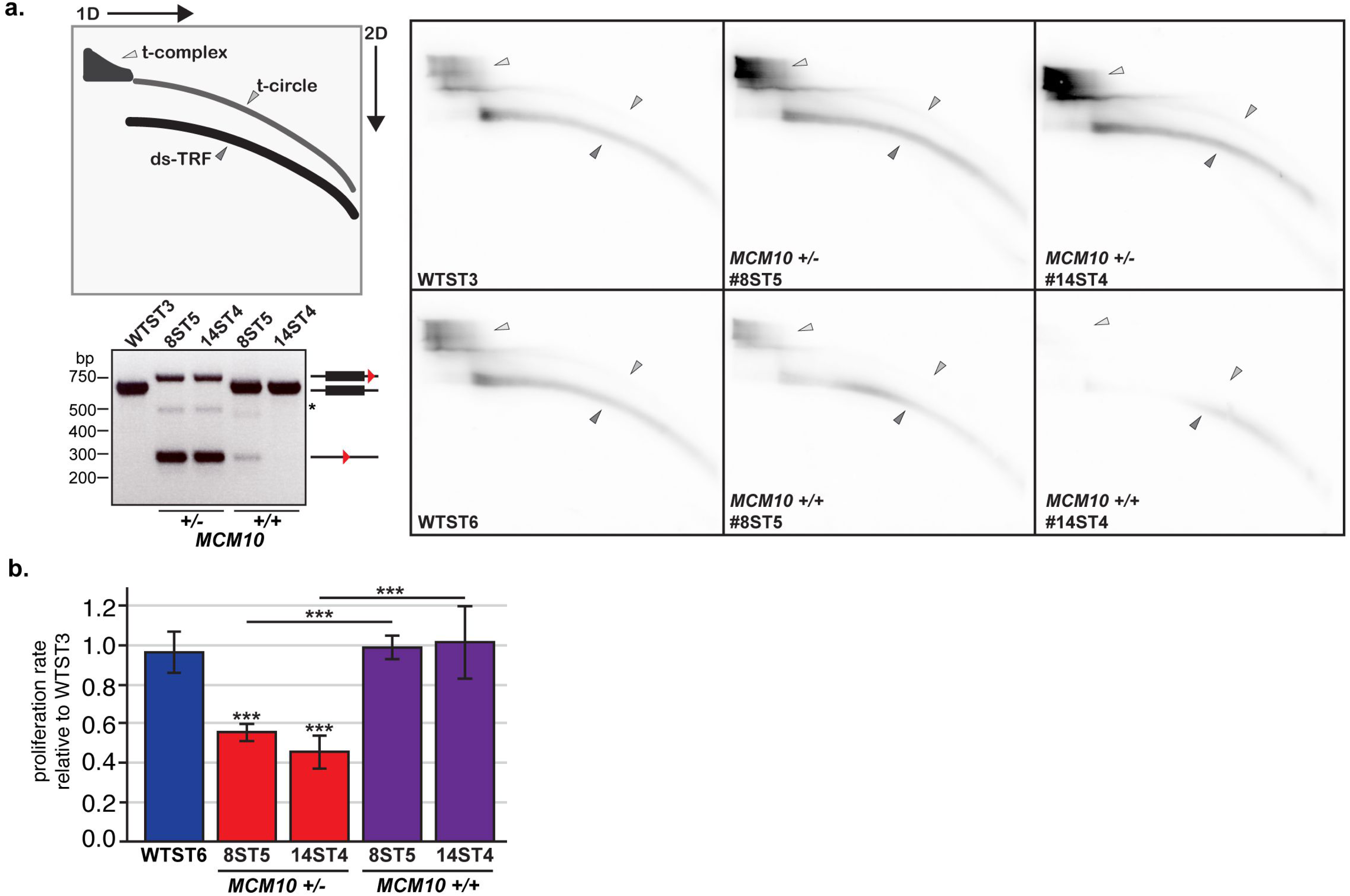
Spontaneous reversion of *MCM10+/-* ST cells rescues t-complex accumulation and proliferation rate. **a)** (Top left) Diagram of double-stranded telomere restriction fragment (ds-TRF), telomere circle (t-circle) and telomere complex (t-complex) DNA species from 2D TRF gel electrophoresis. (Right) Representative comparison of 2D TRF gel electrophoresis in HCT116 wild type, *MCM10^+/-^* and reverted ST cell lines. (Bottom left) Genotyping PCR for exon 14 in cell populations showing alleles that have one *loxP* site 3’ of exon 14 (upper band) or a *loxP* scar (lower band), as well as exon 14 reverted alleles that have retained or lost the 3’ *loxP* site. A faint non-specific band can also be detected (asterisk). **b)** Average proliferation rate in HCT116 wild type, *MCM10^+/-^* and reverted ST cell lines normalized to HCT116 wild type ST3. For each cell line n = 5 replicate wells across two biological replicates; error bars indicate standard deviation and statistical significance was calculated using two-tailed student’s *t-test* with *<.05; **<.01, ***<.001.

**Supplementary table 1.** Overlap of novel chromosomal aberrations in HCT116 wild-type and *MCM10+/-* #8 cell lines with common fragile sites (CFSs).

| HCT116 expected | CFSs from Mrasek et al., 2010 |
| --- | --- |
| dup(10)(q24.1q26.3) | No |
| der(16)t(8;16)(q13;p13.3) | Yes - 8q13 (FRA8F) |
| der(18)t(17;18)(q21;p11.2) | Yes - 17q21 (FRA17D) |
| **Additional aberrations in wild-type (PD ~200)** |  |
| inv(16)(p11.2q23) | Yes - 16q23.2 (FRA16D) |
| der(3)t(3;14)(q27;q11.2) | Yes - 3q27 (FRA3C), 14q11.1 (FRA14D) |
| **Additional aberrations in *MCM10*<sup>+/-</sup> #8 (PD ~25)** |  |
| t(5;8)(p15;q13) | Yes - 5p15 (FRA5H); 8q13 (FRA8F) |
| i(21)(q10) | No |
| der(1)t(1;10)(q21;q11.2) | Yes - 1q21 (FRA1F); 10q11.2 (FRA10G) |
| der(10)t(1;10)(q21;q11.2) | Yes - 1q21 (FRA1F); 10q11.2 (FRA10G) |
| der(5)t(5;10)(p15;q22) | Yes - 5p15 (FRA5H); 10q22.1 (FRAD10D) |
| der(10)t(5;10)dup(10) | No |
| del(11)(p11.2) | Yes - 11p11.2 (FRA11L) |
| del(11)(q23) | Yes - 11q23.3 (FRA11G) |
| dup(7)(q22q32) | Yes - 7q22 (FRA7F), 7q32.3 (FRA7H) |
| del(13)(q22q34) | Yes - 13q22 (FRA13E), 13q31 (FRA13H), 13q32 (FRA13D), 13q34 (FRA13I) |
| del(1)(q32q42) | Yes - 1q32 (FRA1Q), 1q41 (FRA1R), 1q42 (FRA1H) |
| add(15)(p11.2) | No |
| **Additional aberrations in *MCM10*<sup>+/-</sup> #8 (PD ~100)** |  |
| del(6)(q16) | Yes - 6q16.3 (FRA6J) |
| i(15)(q10) | No |
| del(2)(q23) | Yes - 2q23 (FRA2S) |
| add(10)(p11.2) | Yes - 10p11.2 (FRA10J) |
| del(11)(p11.2) | Yes - 11p11.2 (FRA11L) |
| add(3)(q21) | Yes - 3q21 (FRA3M) |
| del(5)(q13) | Yes - 5q13 (FRA5K) |
| del(11)(q13) | Yes - 11q13.3 (FRA11H) |
| add(12)(p11.2) | Yes - 12p11.2 (FRA12H) |
| idic(6)(p23~25) | Yes - 6p23 (FRA6A), 6p25.1 (FRA6B) |
| del(2)(q31) | Yes - 2q31 (FRA2G) |
| idic(15)p12 | No |
| del(3)(p11.2) | No |
| add(19)(p13) | Yes - 19p13.1 (FRA19B) |
| dic(18;22)(p11.2;p11.2) | No |
| add(17)(q25p11.2) | Yes - 17q25 (FRA17E), 17p11 (FRA17C) |
| dic(14;15)(q32;q23~26) | Yes - 14q32 (FRA14H), 15q24 (FRA15), 15q25 (FRA15F), 15q26 (FRA15G) |
| add(17)(q25) | Yes - 17q25 (FRA17E) |
| del(13)(q32) | Yes - 13q32 (FRA13D) |
| t(14;15)(q22;p11) | Yes - 14q22 (FRA14F) |
| add(20)(p11.2) | Yes - 20p11.23 (FRA20A) |
| inv(4)(p14;q31) | Yes - 4p14 (FRA4G), 4q31.1 (FRA4C) |
| der(5)t(5;8)del(5)(q31) | Yes - 5q31.1 (FRA5C) |
| del(6)(q16) | Yes - 6q16.3 (FRA6J) |
| **Overlap with of *MCM10*<sup>+/-</sup> aberrations with CFSs** | **80% (28/35)** |

List of expected and novel chromosomal aberrations identified in HCT116 wild-type or *MCM10* mutant karyotypes.

## Notes

### Competing Interest Statement

The authors have declared no competing interest.

### Summary of Updates

A preprint version of this manuscript was uploaded to bioRxiv previously. At that point, we only described the molecular analysis of MCM10 patient mutations associated with Natural Killer cell deficiency. Our updated report expands the scope of known pathologies caused by defects in replication factors, and argues that specific cell lineages have different requirements for MCM10 expression.

